# DOCK8 regulates a mechanosensitive actin redistribution that maintains immune cell cohesion and protects the nucleus during migration

**DOI:** 10.1101/2024.07.26.605273

**Authors:** Connie Shen, Jérémy Postat, Aysha Cerf, Mauricio Merino, Aanya Bhagrath, Dhanesh Patel, Vincent Luo, Afnan Abu-Thuraia, Caitlin Schneider, Abhinav Sharma, Woong-Kyung Suh, Allen Ehrlicher, Jean-François Côté, Judith N. Mandl

## Abstract

Immune cells navigate through complex 3-dimensional tissue architectures, utilizing an amoeboid mode of migration, characterized by extensive cellular deformation, low adhesion, and high cell velocities. In the absence of expression of *Dedicator of Cytokinesis 8 (Dock8)*, a gene identified with loss-of-function mutations in immunodeficiency, cells become entangled during migration through dense, confined environments and consequently undergo catastrophic cell rupturing, while migration on 2D surfaces remains entirely intact. Here we investigated the specific cytoskeletal defect of *Dock8*-deficient activated T cells, showing that even prior to entanglement they display a striking difference in F-actin distribution compared to wild type (WT) cells. We describe a central pool of F-actin in WT murine and human T cells which is absent in *Dock8* KO T cells, and determine that the relocalization of F-actin is a mechanoresponsive circuit, emerging only when cells are very confined. Our works shows that the central actin pool is nucleo-protective, reducing nuclear deformation and DNA damage during confined migration. We identify the Hippo-pathway kinase Mst1 as a co-mediator of this mechanosensitive pathway in conjunction with Dock8, allowing for cell cohesion and survival during migration through complex environments.

## Introduction

Cell migration is a cornerstone of any immune response. To exert their effector functions and participate in pathogen clearance or tissue homeostasis, leukocytes must be able to traffic throughout the body, transmigrate out of vessels, and traverse diverse tissues in the body which span many distinct architectures, topologies, and mechanical properties^1, 2^. How immune cells are able to navigate such a broad range of 3-dimensional (3D) microenvironments – from lymph nodes, to collagen-rich skin, or dense tumours – and the specific cytoskeletal processes that enable them to do so while keeping their cell shape integrity intact, remains incompletely understood.

Studies of cells migrating in 2D on coated glass *in vitro* have been essential in characterizing the fundamental principles which underlie cell motility^3^. Such 2D studies have described the protrusive filamentous (F-)actin polymerization at the cell front that is driven by the Rho-GTPases Cdc42 and Rac1 and is coupled to actomyosin contractility in the cell rear dominated by RhoA and myosin-II^3–5^. In this setting, integrin-mediated focal adhesions enable cells to pull themselves forward upon polarization^2, 3^. In contrast, in complex 3D environments encountered *in vivo,* leukocytes use an ‘amoeboid’ migration mode where the reliance on, and balance of, adhesive forces required for migration is much more context dependent^6–8^. When migrating in tissue, leukocyte motion often does not require integrins; instead, cells utilize rapid shape changes to push against tissue topology, generating traction forces with little to no tissue remodelling^9–12^. Notably, such pushing forces necessitate an interaction with, and response to, the surrounding substrates, including extracellular matrix (ECM) structures, presumably through mechanically-sensitive pathways^13, 14^. Yet, whether and how cell mechano-responses might enable leukocyte shape adaptations during migration even while moving rapidly through highly confined tissue terrains is still unclear.

Mechanical forces can be sensed and responded to by cells through multiple pathways, including through mechanosensitive ion channels, adhesion molecules, the Hippo pathway transcriptional regulators Yap/Taz, and direct effects on the cytoskeleton, which can in turn modify cell state and behavior^15, 16^. A major limiting factor for the shape changes required during immune cell migration, and of key importance to cell proprioception, is the nucleus, which is the stiffest and bulkiest organelle in the cell^17, 18^. Cells require contractile forces to push the nucleus through small pores^19^, and use their nuclei as a mechanical gauge, allowing the cell to select the ‘path of least resistance’^20^. This nuclear path-sensing is one way in which immune cells can integrate mechanical information about the environment into effective locomotion. In addition, nuclear envelope deformation triggers actomyosin contractility via calcium dependent activation of the phospholipase cPLA2 which leads to arachidonic acid release^21, 22^. Overall, the physical confinement experienced by immune cells when embedded in tissues, together with the extreme shape deformations they have to undergo to squeeze past obstacles, imposes challenges on the cell that may require additional or alternate molecular pathways to maintain their cohesion that are dispensable in 2D.

Inherited defects in immune cell migration provide an opportunity to probe the specific pathways required in navigating 3D tissue environments. Such is the case with the loss of expression of *dedicator of cytokinesis 8* (Dock8), which we and others have previously described to be critical for cell shape integrity during migration in highly confined 3D environments for T cells and dendritic cells (DCs)^23–25^. *Dock8* KO cells display dysregulated morphology, loss of cell cohesion, and ultimately cell death by migration-induced shattering, termed ‘cytothripsis’^24^. Dock8 is an atypical guanine exchange factor (GEF) expressed only in immune cells which binds and regulates Rho GTPases by facilitating the exchange of GDP for GTP^26^. Dock8 belongs to the 11-member family of DOCK180-related GEFs, a family that can interact with GTPases such as Rac, RhoA, and Cdc42, which are important for actin and cytoskeletal rearrangements implicated in cellular functions such as cell motility, growth, survival, polarity, and differentiation^27, 28^. Rho GTPases modulate actin polymerization through actin binding proteins such as Arp2/3, via activation of nucleation promoting factors such as WASP and WAVE^29–31^. Importantly, failure of *Dock8-*deficient leukocytes to migrate effectively in tissues leads to an immunodeficiency clinically characterized by disseminated cutaneous and systemic infections, hyper IgE syndrome, and allergic disease^32–34^.

Here, we employed *in vitro* migration assays to understand the molecular underpinnings of how Dock8 regulates cell cohesion and nuclear integrity in T cells. We found that in the absence of environmental complexity, *Dock8*-deficient T cells display no detectable cytoskeletal defects, and in fact migrate at greater speeds and pass through narrow pores with greater ease than wild type (WT) T cells. However, in confined conditions, Dock8 is required for a redistribution of F-actin from the cell front to the cell center near the nucleus. Our data suggest that this central F-actin pool protects the nucleus, prevents force-mediated DNA damage, and is key to balancing the forward propulsion of the cell’s leading edge with the necessity to maintain cohesion as the cell becomes stretched in confined spaces. We determined that this mechanosensitive actin redistribution response is dependent on both Dock8 and the Hippo-pathway kinase Mst1. Together, our data indicate that Dock8 is therefore a critical mediator of leukocyte mechanosensing during migration in 3D environments.

## Results

### Loss of cell cohesion and consequent death of migrating *Dock8* KO T cells is collagen density-dependent

To investigate the impact of *Dock8*-deficiency on immune cell migration dynamics in 3D, we focused on activated T cells, which upon antigen encounter *in vivo* gain access to a variety of non-lymphoid tissue landscapes to curtail pathogen spread^35^. As one of the softest immune cell types^36^, T cells have the challenging task to maintain cell cohesion even as they deform to traverse tight gaps in the ECM or between other cells. We activated WT or *Dock8* KO murine T cells *in vitro* using anti-CD3/CD28 for 4 days and then embedded them in bovine collagen gels, previously used to mimic complex 3D tissue structure *in vitro*^37^, and in which T cells move spontaneously without the addition of chemokine. Characteristic of their amoeboid migration in these gels, WT T cells frequently and rapidly modulated their shape to navigate past collagen fibers, but *Dock8* KO T cells displayed much greater shape distortions, including extreme elongations (**Fig. 1a** and **Video S1**). Such cell elongation and entanglement of *Dock8* KO T cells was not necessarily permanent in all instances, as some cells recovered to resume motility (**Fig. 1a**, bottom example). Notably, as shown previously, the addition of CCL19 to one side of the collagen gel to generate a gradient showed that chemotaxis by *Dock8* KO T cells in 3D collagen remained intact, although cell speed was slightly decreased, as was track straightness (**Fig. S1a-c**).

**Figure 1.**
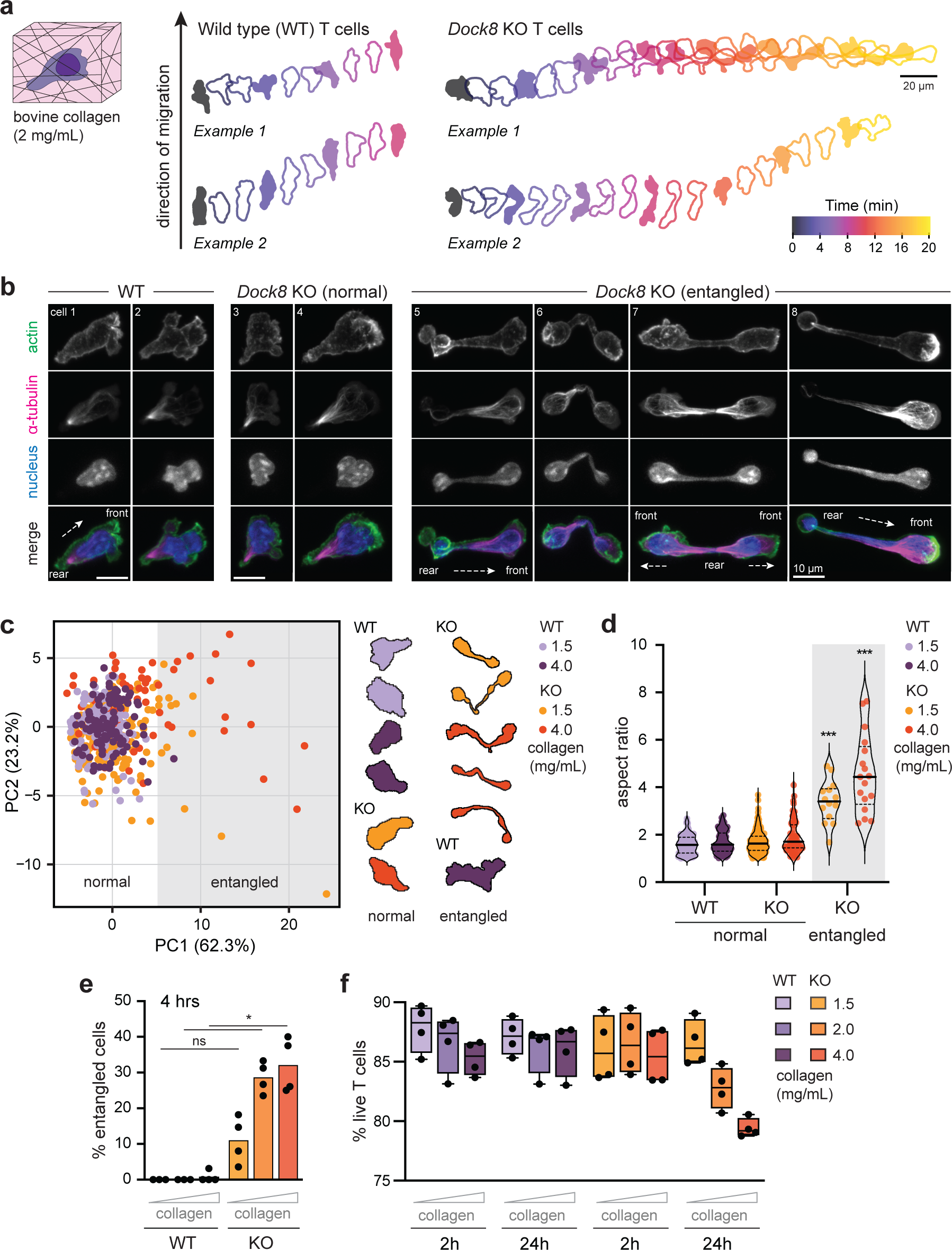
Loss of cell cohesion and death of migrating *Dock8* KO T cells is collagen-density dependent. (a) Two representative cell outlines of activated WT and *Dock8* KO T cells spontaneously migrating through a collagen matrix made of 2 mg/mL bovine type I collagen (no chemokine added), colour-coded by time with the direction of migration indicated (arrow). **(b)** Representative confocal microscopy images (max projection of z-stacks) of T cells migrating through collagen matrix as in (a). Examples of WT and *Dock8* KO T cells (entangled or not) are shown with cell front and rear indicated. Fixed cells were stained for F-actin (phalloidin), tubulin (α-tubulin antibody), and nucleus (Hoechst). **(c)** Principal component analysis (PCA) of 20 extracted cell shape parameters from *n =* 211 WT and *n =* 293 KO T cells migrating in low and high collagen densities (1.5 and 4 mg/mL) after 4 hours. Data points are individual cells; example cell outlines of T cells classified as normal or entangled by PCA are shown. **(d)** Summary violin plots of the aspect ratio of cells analysed in (c) for each category. Data points: individual cells; solid lines: median; dotted lines: quartiles. **(e)** Summary plot of percent WT and *Dock8* KO T cells entangled in collagen matrix (2 mg/mL) after 4 hours of migration. Data points: replicate T cell cultures from *n =* 4 mice; means are indicated. **(f)** Summary plot of percent live WT and *Dock8* KO T cells in collagen matrix (1.5, 2, or 4 mg/mL) after 2 or 24 hours of migration. Data points: replicate T cell cultures from *n =* 4 mice; medians are indicated with quartiles (box) and min to max ranges (error bars). Statistical tests: Kruskal-Wallis ANOVA with Dunn’s multiple comparison test (d, e). \**P <* 0.05; \*\*\**P <* 0.001; ns, non-significant.

Next, we examined microtubule and F-actin arrangement in fixed WT and *Dock8* KO T cells migrating in collagen. Motile WT T cells had a polarized morphology with the microtubule organizing center (MTOC) in the uropod, the actin-rich cell front probing into the environment, and the nucleus deforming to fit through the constraints imposed by the collagen matrix (**Fig. 1b**, examples 1 and 2). In *Dock8* KO T cells migrating normally, the gross morphology observed in WT T cells was conserved, with no defect in cell polarization (**Fig. 1b**, examples 3 and 4). However, in entangled *Dock8* KO T cells, both the tubulin and F-actin organization was substantially dysregulated, with the nucleus stretched throughout the entire cell body. Most commonly, entangled *Dock8* KO cells had two competing cellular fronts at opposite ends, with more diffuse tubulin throughout the cell, and a centrally located MTOC (**Fig. 1b**, examples 6 and 7, and **S1d**). Less commonly, *Dock8* KO cell rears were ‘stuck’ in the matrix with the cell front continuing to advance (**Fig. 1b**, example 5 and 8). Overall, entangled *Dock8* KO T cells were exclusively bipolar (**Video S2**), different to what was observed in *Dock8* KO DCs^23^.

Importantly, the collagen-rich and highly cross-linked ECM structure of the skin may explain the inability of *Dock8* KO T cells to control skin viral and bacterial infections in particular, hence leading to the characteristic cutaneous manifestations of Dock8-deficiency^24, 38^. Thus, we next explored the relationship between collagen density and the dependence of T cells on *Dock8* expression for cell shape integrity in 3D by comparing cell morphology in 1.5, 2 or 4 mg/ml collagen which differed in fiber density, pore size, and scaffold stiffness (**Fig. S1e,f**)^39^. To systematically quantify cell shape integrity in WT and *Dock8* KO T cells at low or high collagen concentrations, we fixed cells after 4 hours of migration when the majority of both WT and *Dock8* KO cells were still viable, extracted cell shape outlines which we parameterized with 20 cell morphology descriptors (**Table S1**), and performed a principal component analysis (PCA). We found that entangled *Dock8* KO cells reliably segregated from the majority of cells along PC1 based on cell shape alone (**Fig. 1c**). The cell aspect ratio (the length of the major cell axis divided by the minor axis) was one of the best predictors of cell entanglement (increased by ∼3-fold in *Dock8* KO cells that had lost their shape integrity), corroborating the consistently elongated shape characteristic of entangled T cells (**Fig. 1b**). While staining for F-actin showed that WT T cells became increasingly constrained as the collagen concentration increased, most evident from the increasingly irregularly shaped cell front as cells were probing tighter paths, they nonetheless maintained shape integrity (**Fig. S1g**). In contrast, *Dock8* KO T cells became entangled at all collagen densities tested, but the fraction of entangled cells increased with greater collagen density (**Fig. 1e** and **S1g**). Moreover, *Dock8* KO T cell viability after 24 hours of migrating in collagen decreased as collagen density was increased, correlating with the fraction of cells that lost cohesion (**Fig. 1f**). Together, our data showed that *Dock8* KO T cells have a defect in 3D migration whereby cells became increasingly entangled, lost cell shape integrity, and died as collagen concentration, and thus environmental complexity, was increased.

### *Dock8* KO T cells navigate simple environments and tight constrictions with no impairment

To better understand the migration defect observed in *Dock8* KO T cells, we next investigated their motility in simplified environments where we could precisely control the specific constraints or obstacles encountered (without chemokine added). For this, we turned to fibronectin-coated microfluidic devices fabricated from polydimethylsiloxane (PDMS) (**Fig. 2a**)^40^. First, we characterized cell migration speeds in one-dimensional straight channels wide and high enough (6mm x 5mm) for activated T cells to traverse easily without deformation. Surprisingly, we found that *Dock8* KO T cells were considerably (∼1.6 fold) faster on average than WT T cells (**Fig. 2b,c** and **Video S3**). The greater migration speed of *Dock8* KO T cells was maintained in pillar forests (∼1.5 fold fast than WT T cells), in which cells had to navigate obstacles and bifurcating paths but channel widths and heights were the same as for the straight channels (**Fig. 2d,e**). Notably, we did not observe the loss of cell cohesion of *Dock8* KO T cells in the simple lattice structure of the pillar forests (**Fig. 2e** and **Video S4**).

**Figure 2.**
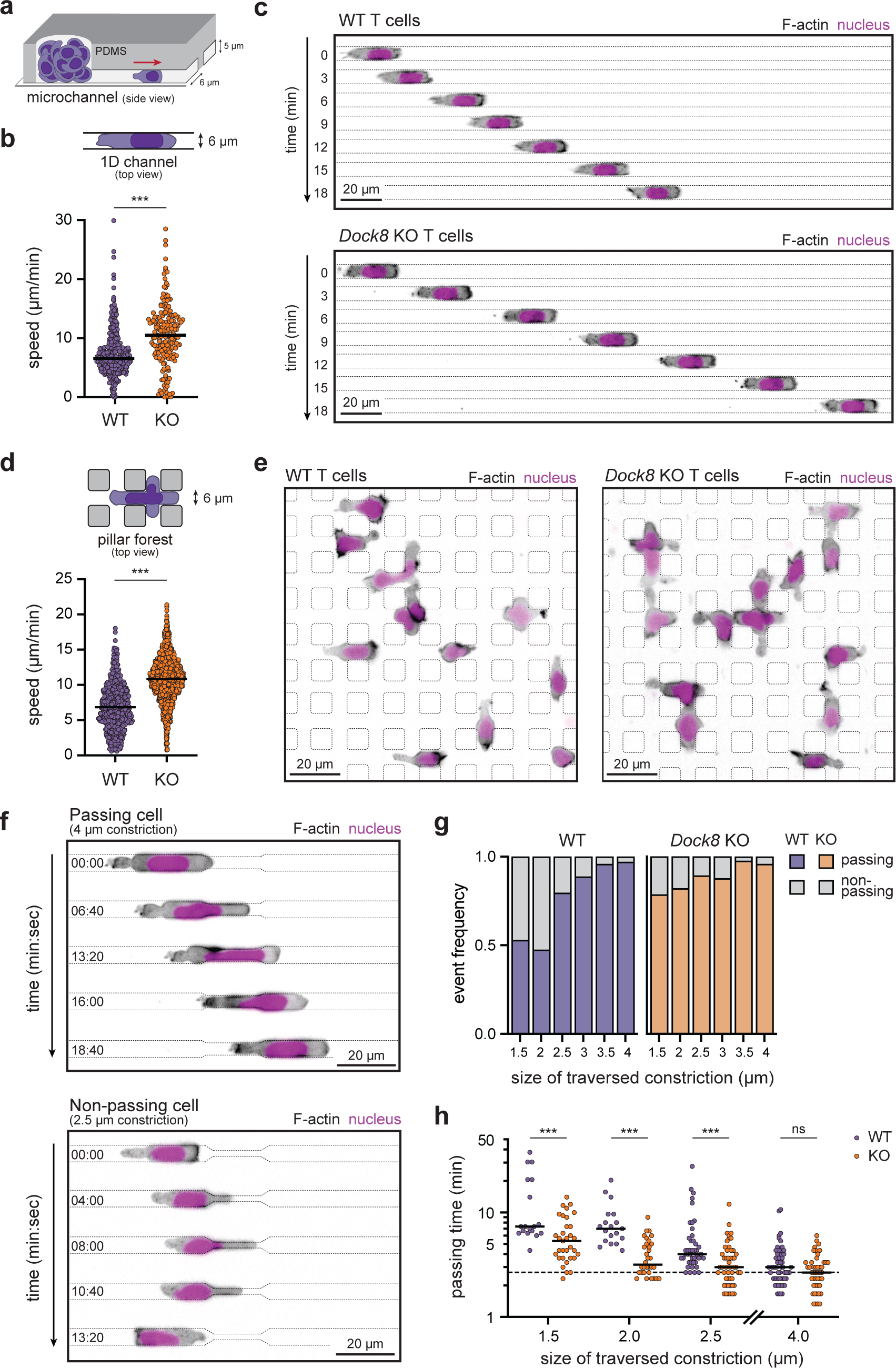
*Dock8* KO T cells navigate simple environments and small constrictions with no impairment. (a, b, c) Activated WT and *Dock8* KO T cells expressing LifeAct-GFP and NLS- nTnG migrating in straight microchannels (width of 6 µm and height of 5 µm) coated with fibronectin, no chemokine added. Summary of individual cell mean speed (b), and representative examples acquired by epifluorescence microscope shown over time (c). Data is from 2 independent experiments, *n =* 361 (WT) and *n =* 179 (KO) cells; line: median. **(d, e)** Activated WT and *Dock8* KO T cells expressing LifeAct-GFP and NLS-nTnG migrating through pillar forest microchannels (channel width of 6 µm, height of 5 µm, pillars of 8 µm x 8 µm) coated with fibronectin, no chemokine added. Summary of individual cell mean velocities (d), and representative examples acquired by epifluorescence microscope (e). Data is *n =* 572 (WT) and *n =* 1385 (KO) cells; line: median. **(f, g, h)** Activated WT and *Dock8* KO T cells expressing LifeAct-GFP and NLS-nTnG migrating through microchannels with constrictions (channel width of 6 µm, height of 5 µm, constriction sizes 1.5–4 µm) coated with fibronectin, no chemokine added. Representative examples of cell retreating and cell passing acquired by epifluorescence microscope (f). Percent of cells passing through or retreating at constrictions (g), and summary of time spent passing through constrictions of different sizes, data points are individual cells, *n =* 18–57 cells per constriction size; line: median; dotted line at average passing time through 4 µm constrictions (h). Statistical tests: Mann-Whitney test (b, d); 2-way ANOVA (h). \*\**P* < 0.01; \*\*\**P <* 0.001; ns, non-significant.

As the loss of cell shape integrity of *Dock8* KO T cells in collagen matrices appeared to be triggered by becoming stuck in the collagen fiber mesh, we next asked whether *Dock8* KO T cells had difficulty traversing narrow constrictions. To address this, we used straight channels as before, but incorporated constrictions 15 µm in length, ranging from 1.5 to 4 µm in width. Surprisingly, we found that *Dock8* KO T cells successfully passed through even the smallest constrictions at a greater frequency compared to WT T cells, and of T cells able to pass through constrictions, the passage time of *Dock8* KO cells was consistently shorter than that of WT counterparts (**Fig. 2f-h** and **Video S5**). A rate-limiting step in the ability of cells to squeeze through tight spaces is the rigidity of the nucleus^18^, the largest organelle in the cell, and transient nuclear envelope rupture has been shown to occur in some cell types to facilitate passage of constrictions^41, 42^. Thus, the greater ease with which *Dock8* KO T cells traversed narrow constrictions could indicate either that *Dock8* KO T cells (and their nuclei) are inherently more deformable, or that their nuclear envelope is more prone to rupture, enabling passage through constrictions. Our observation that the nucleus in entangled *Dock8* KO cells was stretched could be compatible with either explanation, given that we used the DNA intercalating agent Hoechst to visualize the nucleus in collagen (**Fig. 1b**). To date, it has not been investigated whether nuclear envelope rupture could impact T cell migration through small pores. To probe this further, we isolated T cells from mice expressing tdTomato fluorescent protein with a nuclear localization signal (NLS-nTnG) and examined whether *Dock8* KO T cells underwent nuclear rupture when passing through constrictions. We found that in both WT and *Dock8* KO T cells passing through even the smallest 1.5 µm widths, nuclear envelope rupture was a rare occurrence, accounting for fewer than 1% of all observed passing events (data not shown). When nuclear envelope ruptures did occur, the fluorescent reporter leaked into the cell body and could be detected in the cell cytoplasm (**Fig. S2a** and **Video S6**). Similarly, we found that although the nucleus of *Dock8* KO T cells entangled in collagen matrices lost its shape integrity, the nuclear envelope remained intact (**Fig. S2b** and **Video S7**). These results indicate that *Dock8* KO T cells are inherently faster than WT T cells when migrating in simple environments, and that KO cells have more deformable nuclei without increased susceptibility to nuclear envelope rupture.

### Dock8 is necessary for a mechanoresponsive, integrin-independent redistribution of F-actin

Because we observed that simple environmental obstacles such as pillars or constrictions did not lead to loss of cell shape cohesion in *Dock8* KO T cells, we next asked whether confinement in the absence of obstacles would present a challenge, and whether a dysregulation of the organization of intracellular compartments or organelles in *Dock8* KO T cells might play a role in the migration defect. To do so, we utilized an under-agarose assay, where cells migrated freely in 2D, compressed between a fibronectin-coated glass slide and a pad of agarose, towards the chemokine CCL19^43^. We found no difference in the arrangement or levels of acetylated tubulin or α-tubulin, the location of the MTOC, levels or location of G-actin, mitochondria, lysosomes or vimentin between migrating WT or *Dock8* KO T cells (**Fig. S3a-e)**. However, we observed a striking difference in the localization of F-actin stained in fixed cells (using phalloidin) between confined WT and *Dock8* KO T cells (**Fig. 3a**). Without confinement, both WT and *Dock8* KO T cells migrating in 2D had the expected enrichment of F-actin at the leading edge with little actin polymerization occurring at the cell centroid (**Fig. 3a,b**). In contrast, in confined WT T cells, a distinct pool of F-actin appeared in the cell center which was entirely absent across all *Dock8* KO T cells analysed (**Fig. 3a,b**). This difference in F-actin localization reflected a change in distribution, as the total F-actin levels were similar between WT and *Dock8* KO T cells (**Fig. S3f,g**). Notably, when *Dock8* KO T cells encountered other cells under agarose, this was sufficient to lead to an entangled phenotype as observed in the collagen gels (**Fig. S3h** and **Fig. 1**). The phenotype of cells with a loss in shape integrity had a quite distinct organization from *Dock8* KO T cells undergoing cell division (**Fig. S3i**).

**Figure 3.**
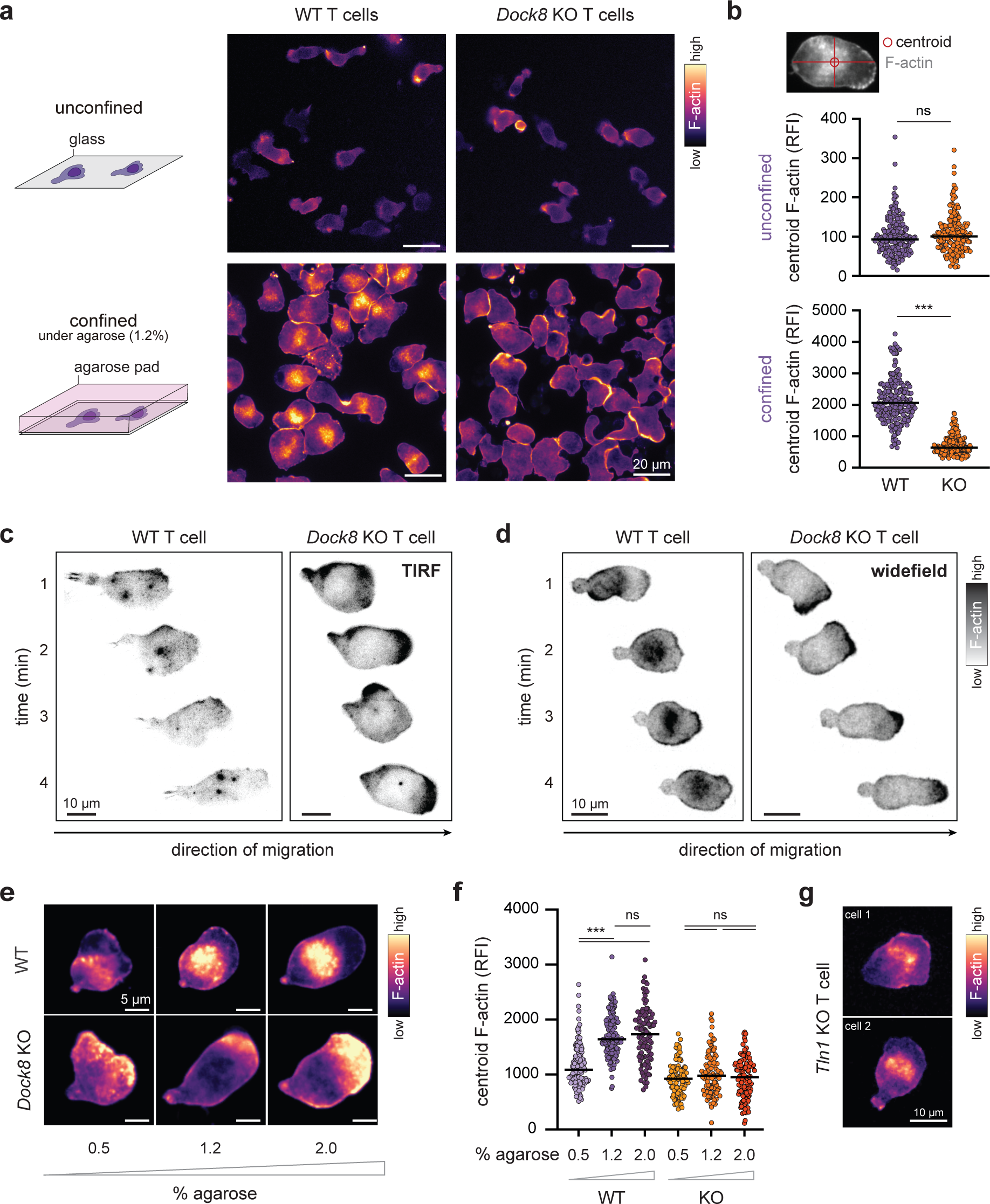
Confinement-induced redistribution of F-actin is absent in *Dock8* KO T cells. (a) Fluorescence intensity of F-actin (phalloidin) in WT or Dock8 KO T cells migrating on fibronectin-coated glass either in absence of (unconfined) or under 1.2% agarose (confined). Representative images (epifluorescence microscope) are shown. **(b)** Summary plot of F-actin relative fluorescence intensity (RFI) measured at the cell centroid as indicated (top). Data points, individual cells (*n =* 166-210 cells); lines, medians. **(c, d)** Representative WT and *Dock8* KO T cells expressing LifeAct-GFP (grey scale, fluorescence intensity) migrating under 1.2% agarose over time acquired by TIRF (c), or widefield microscopy (d). **(e, f)** Fluorescence intensity of F- actin (phalloidin) in WT and *Dock8* KO T cells migrating under 0.5, 1.2 or 2% agarose. Representative examples (epifluorescence microscope) (e), and summary plots (f). Data points, individual cells (*n =* 109-140 cells); lines, medians. **(g)** Two representative example images of F- actin fluorescence intensity in talin1 (*Tln1*)-deficient murine T cells migrating under 1.2% agarose. Statistical tests: Mann-Whitney tests (b); Kruskal-Wallis ANOVA with Dunn’s multiple comparison test (f). \*\*\**P <* 0.001; ns, non-significant.

To further characterize the confinement-dependent central F-actin pool, we tracked the dynamics of cortical actin in direct contact with the fibronectin-coated glass in LifeAct-GFP expressing T cells, using total internal reflection fluorescence (TIRF) microscopy. In WT T cells we observed the transient appearance of many small actin patches throughout the cell, which existed on the time scale of seconds (**Fig. 3c**), as were noted also in confined DCs^13^. In *Dock8* KO T cells, these transient actin patches were reduced in number, and the majority of F-actin was distributed more uniformly at the cell front (**Fig. 3c** and **Video S8**). Examining the entirety of LifeAct-GFP T cells by widefield microscopy, we again observed a clear F-actin pool in the center of the WT T cells that, while dynamic in shape and intensity, was stable in its location at the mid- zone of the cell, and which was completely absent in *Dock8* KO T cells (**Fig. 3d** and **Video S9**). As the appearance of the central F-actin cloud was only seen under confinement, we next asked whether this mechanosensitive F-actin response would be modulated by the rigidity of confining agar. To address this, we titrated the percent agarose from 0.5 to 2% to increase the degree of mechanical load^44^. Indeed, the fluorescence intensity at the cell centroid of F-actin scaled with the percent agarose in WT T cells (**Fig. 3e,f**). Moreover, even at the highest agarose percent and thus the greatest level of confinement, *Dock8* KO T cells did not redistribute F-actin to the cell center (**Fig. 3e,f**).

Although dispensable for T cell motility under confinement^9^, integrins are a well-described mediator of mechanotransduction^16^, and in NK cells Dock8 was found to be part of a multi-protein complex that included talin1, a required cytosolic adaptor protein for integrin-mediated signaling^45^. Thus, we next tested whether the mechanosensitive F-actin rearrangement in WT T cells was dependent on integrin signaling by confining talin1-deficient (*Tln1* KO) T cells under agarose. *Tln1* KO T cells were previously shown to be unable to adhere to 2D surfaces^9^. We found that *Tln1* KO T cells remained similarly able to rearrange the F-actin to the cell center in response to mechanical force as WT T cells, indicating that the appearance of the mechanosensitive F-actin pool did not require integrin signaling (**Fig. 3g**). These data indicated that T cells redistribute cellular actin in direct response to mechanical load and that this mechanosensitive cytoskeletal response requires Dock8 expression but is integrin signaling-independent.

### The confinement-induced central actin pool is located at the nucleus front in human and murine T cells

We next sought to understand whether the central F-actin pool was located in a specific region of the cell, particularly in relation to cellular organelles. We hypothesized that its appearance in the cell center might be tied to the location of the nucleus. T cells migrating under-agarose were fixed and stained with both phalloidin and Hoechst, to examine the distribution of F-actin per cell from rear to front. One example T cell of each of WT and *Dock8* KO (**Fig. 4a,b**), as well as across n = 50 cells per genotype (**Fig. 4c**) were analysed. While in confined migrating WT T cells there was a consistent peak in F-actin fluorescence intensity located towards the front of the nucleus, in *Dock8* KO T cells the fluorescence intensity of F-actin was lowest in proximity to the nucleus (**Fig. 4a-c**). Moreover, during migration under confinement, the F-actin cloud in WT T cells, while dynamic alongside cellular shape changes, maintained its position towards the nucleus front in the direction of migration (**Fig. 4d** and **Video S10**). Importantly, we confirmed that the presence and positioning of the confinement-dependent central F-actin pool was similar for migrating activated human T cells (**Fig. 4e**).

**Figure 4.**
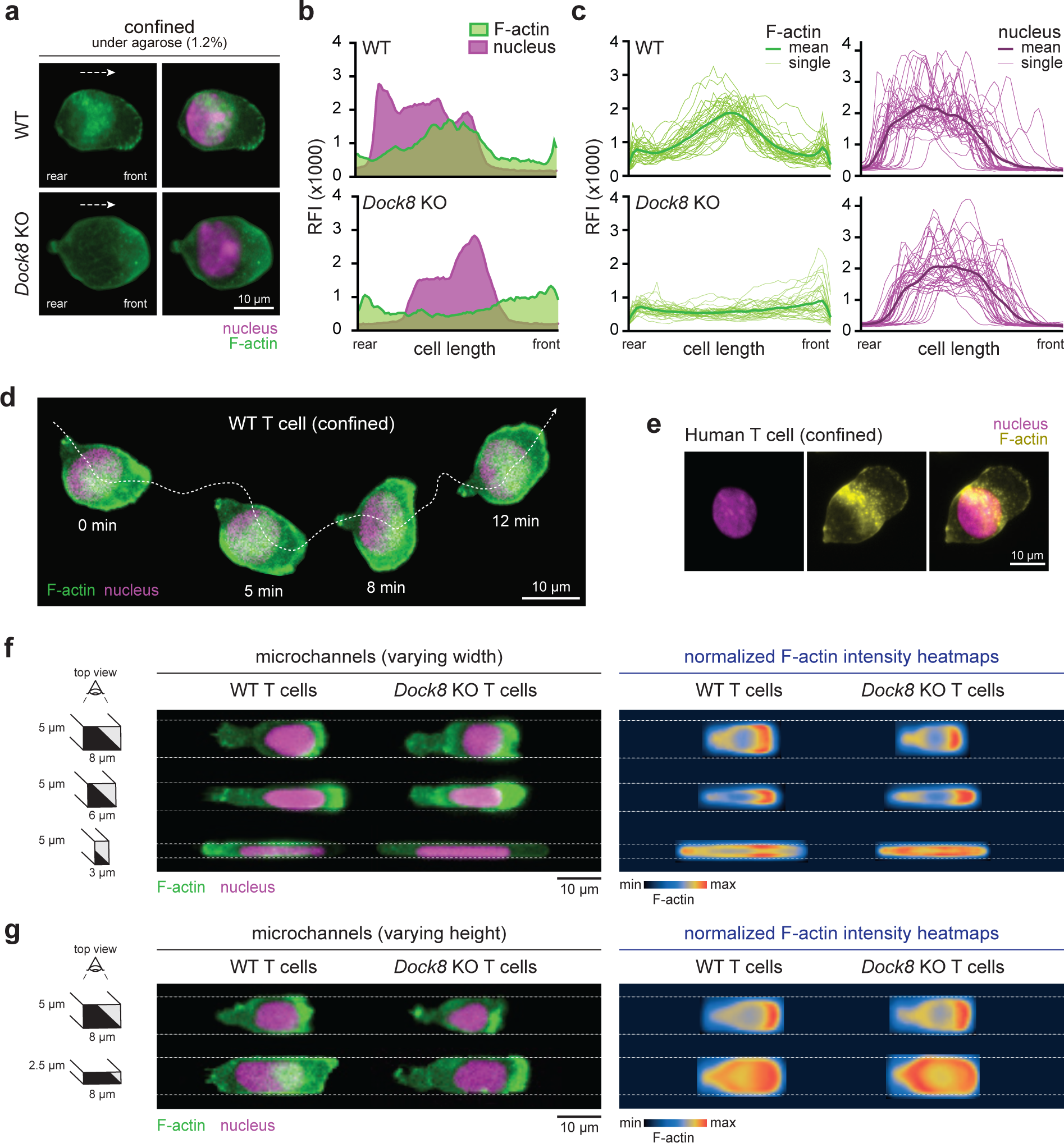
The confinement-induced central actin pool is located at the nucleus front in migrating human and murine T cells. (a) Representative images of WT and *Dock8* KO T cells migrating under 1.2% agarose. Cells were fixed and stained for F-actin (phalloidin) and nucleus (Hoechst), and images were acquired by epifluorescence microscopy. **(b)** Histograms of RFI for both F-actin and nucleus from cells shown in (a) taken cross-sectionally across the cell from rear to front. **(c)** Aggregated histograms of F-actin (phalloidin) and nucleus (Hoechst) RFI of individual T cells (*n =* 35 per genotype) normalized over cell length by linear interpolation. Lines for individual cells and means are shown. **(d)** Representative timelapse images of WT T cells expressing LifeAct-GFP and NLS-nTnG migrating under 1.2% agarose acquired by epifluorescence microscopy. **(e)** Representative human T cell migrating under 1.2% agarose, fixed and stained for F-actin (phalloidin) and nucleus (Hoechst). **(f, g)** Representative images (left) and normalized F-actin intensity heatmaps of *n =* 12-31 cells (right) of WT and *Dock 8* KO T cells expressing LifeAct-GFP and NLS-nTnG migrating in straight microchannels with either fixed heights of 5 µm and variable widths of 3, 6 or 8 µm (f), or variable heights of 2.5 or 5 µm and fixed widths of 8 µm (g).

Our data thus far suggest that the intensity of the F-actin pool in WT T cells scaled with greater mechanical force imposed by increasing agarose concentrations. To better define the extent of confinement required to elicit the cytoskeletal actin rearrangement, we next used microchannels with variable widths from 8µm to 3µm but constant heights of 5µm, or variable heights of 5µm or 2.5µm but constant widths of 8µm. We showed that only when cells were confined in either widths or heights below 3 µm the F-actin redistribution was triggered in WT T cells (**Fig. 4f,g**). We confirmed that also during migration through confined microchannels *Dock8* KO T cells did not relocate F-actin to the nuclear region (**Fig. 4f,g**). Thus, in both murine and human activated WT T cells, migration under confinement triggers a redistribution of polymerizing actin to a nucleus-proximal central region within cells.

### Cytoskeletal response and cell shape cohesion under confinement is Dock8-dependent in dendritic cells, but not neutrophils

Given that we and others have previously described that DCs exhibited migration defects when they lacked Dock8 expression^23, 25^, we next investigated whether we also observed a central F-actin pool in bone marrow derived DCs (BMDC) under confinement, and whether it was similarly Dock8-dependent as in T cells. In an under-agarose migration assay, we found that WT BMDCs had a large increase in centrally located actin polymerization that was absent in *Dock8* KO BMDCs (**Fig. S4a**). Compared to WT T cells, the central actin pool in WT BMDCs occupied a much smaller proportion of the cell (**Fig. S4a,b**). We also examined the F-actin distribution within neutrophils migrating under agarose, and unexpectedly detected no centrally relocated F-actin pool in either WT or *Dock8* KO neutrophils (**Fig. S4c**). In both WT and *Dock8* KO cells, the F-actin distribution in confined neutrophils remained largely localized towards the front of the cell, particularly in the protrusive regions (**Fig. S4c**). The absence of the mechanically-induced central F-actin pool in neutrophils suggested the possibility that, unlike T cells and BMDCs, cell shape cohesion during 3D migration of neutrophils might be independent of Dock8 expression. To test this, we embedded LifeAct-GFP and NLS-nTnG-expressing WT and *Dock8* KO neutrophils in 2 mg/ml collagen and followed their morphology over time. Interestingly, *Dock8* KO neutrophils did not lose cell cohesion, become entangled or abnormally stretched during migration (**Fig. S4d,e**). Taken together, BMDCs and T cells, but not neutrophils, rely on a Dock8-dependent mechanosensitive pathway to redistribute actin towards the center of the cell. Moreover, the loss of the central F-actin in Dock8-deficiency leads to a migration defect in 3D in BMDCs and T cells, but is dispensable for cell shape integrity in confined environments in neutrophils.

### Dock8-dependent F-actin redistribution in migrating T cells is nucleo-protective

Our data from distinct leukocyte types suggested that the redistribution of F-actin was associated with the maintenance of cell cohesion during 3D migration. We next investigated whether the confinement-induced central F-actin pool might have an additional functional role in migrating cells. To do so, we asked whether *Dock8*-deficiency in T cells led to differences in gene expression. We performed RNA sequencing on WT or *Dock8* KO activated CD8^+^ T cells after 24 hours of migration in collagen matrices or incubation in media (non-migrating). Interestingly, we found that the majority of gene expression differences appeared in migrating WT compared to KO cells. Only 19 differentially expressed genes (DEG) were identified in the media condition, one of which was *Dock8* itself, compared to 925 DEG identified in collagen-embedded WT versus KO cells (**Fig. 5a**). Examining the pathways which were most perturbed during migration in *Dock8* KO T cells using gene ontology analysis, we found an enrichment of genes related to cell motility, cell death, adhesion, and actomyosin organization, in line with the observed cytoskeletal defect and migration-induced death of *Dock8* KO T cells (**Fig. 5b**). We also found that a number of genes in the ‘p53 signaling’ pathway, as well as in the ‘response to virus’ pathway were significantly enriched in migrating *Dock8* KO T cells. This suggested the possibility that *Dock8* KO T cells were accruing DNA damage and activating p53 as a result, with associated type I interferon signaling due to a DNA damage-related response downstream of DNA sensors. Indeed, among upregulated genes in *Dock8* KO T cells was *Aim2* (DNA sensor triggering inflammasome activation), *Oasl1, Oasl2,* and *Mx1* (classic type I IFN response genes), as well as p53 target or DNA damage response-related genes such as *Aen, Cdkn1a, Epha2, Phlda3, Pmaip1* (**Fig. 5c**).

**Figure 5.**
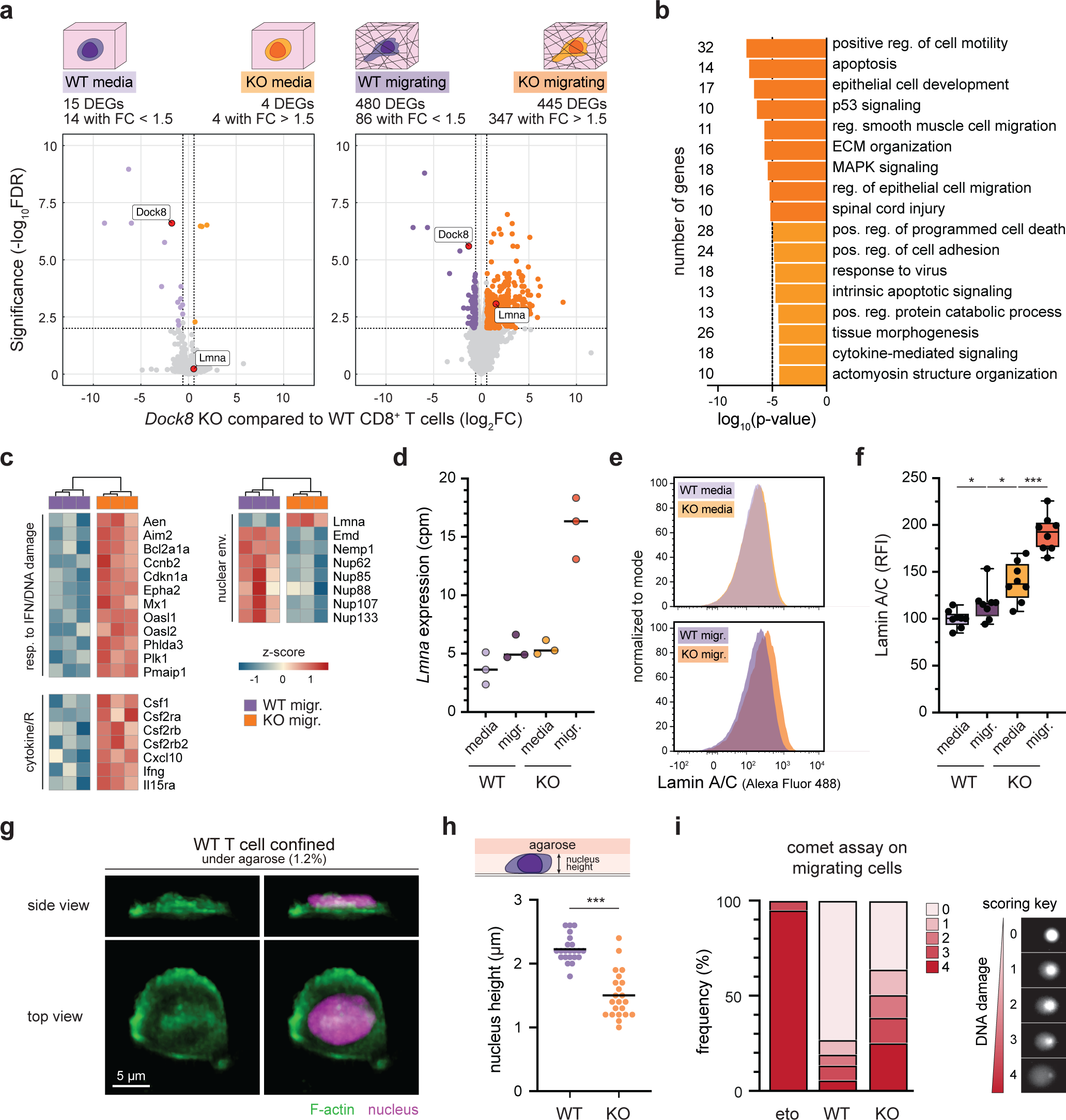
The DOCK8-dependent F-actin redistribution in migrating T cells is nucleo- protective. (a) Volcano plots of significant differentially expressed genes (DEG) identified by RNA-seq between WT and *Dock8* KO CD8^+^ T cells activated *in vitro* and cultured in either media or migrating in collagen (2 mg/ml) for 24 hours with 3 biological replicates per genotype (*FDR* < 0.01). FC, fold change. Specific genes of interest (red data points) are labeled. **(b)** Gene ontology analysis of significant DEGs (445 genes) upregulated in *Dock8* KO compared to WT T cells from (a) at FC > 1.5. Bar graph showing gene numbers per category of the top non- redundant enrichment clusters identified with at least 10 genes. Reg., regulation; pos., positive. **(c)** Heatmap of expression (z-score) of selected genes in WT and *Dock8* KO T cells identified among DEGs from (a). **(d)** *Lmna* gene expression from RNA-seq data. cpm, counts per million. Data points are independent *in vitro* activated CD8^+^ T cell cultures from *n =* 3 mice per genotype; lines represent medians. **(e,f)** Lamin A/C expression quantified by flow cytometry in WT or *Dock8* KO CD8^+^ T cells cultured in either media or migrating in collagen (2 mg/ml) for 24 hours. Representative flow cytometry histograms (g), and data summarized from *n =* 8 mice per genotype and 3 independent experiments (h); lines represent medians; boxes are quartiles; error bars represent the min to max range; RFI are normalized to the mean of WT T cells in media. **(g,h)** LifeAct-GFP and NLS-nTnG WT and *Dock8* KO T cells migrating under 1.2% agarose and imaged using lattice light sheet microscopy. Representative image of a WT T cell shown both from a top view and a side view (g). Summary plot of nuclear height measurements taken from cells that were polarized and not in contact with more than one other cell (h). Data points, *n =* 22 cells per genotype; lines, means. **(i)** Alkaline comet assay of WT T cells treated with etoposide (n = 61 cells), or WT (n = 75 cells) and *Dock8* KO (n = 52 cells) T cells migrating in collagen (2 mg/ml) for 12 hours. Representative images of visual scoring of double- stranded DNA breaks (score 1 – 4) are shown (right). Statistical tests: one-way ANOVA with Holm-Šídák’s multiple comparisons test (f); Mann-Whitney test (h); \**P* < 0.05; \*\*\**P <* 0.001; ns, non-significant.

In addition to several cytokine or cytokine receptor genes upregulated in migrating *Dock8* KO T cells compared to their WT counterparts, genes encoding nuclear envelope proteins were differentially regulated (**Fig. 5c**). This included *Lmna*, which encodes for lamin A/C – a component of the meshwork of structural fibrous proteins protecting the cell’s genomic material and a key determinant of nuclear stiffness^46, 47^. In our RNA sequencing data, *lmna* expression was comparable between WT T cells in media or collagen and *Dock8* KO T cells in media, but significantly upregulated (∼3 fold) in *Dock8* KO T cells migrating in collagen (**Fig. 5d**). When we measured lamin A/C protein levels by flow cytometry, we found that there was a slightly greater level of lamin A/C in *Dock8* KO T cells compared to WT T cells when kept in media, but that this difference increased between KO and WT T cells migrating in collagen (**Fig. 5e,f**). Nuclear stiffening through the increase of lamin A/C has been shown to be a cellular response to mechanical cues^48, 49^. This suggests that *Dock8* KO T cells might be experiencing greater mechanical force transmitted onto the nucleus than WT T cells during migration as a result of the loss of the central, nucleus-proximal F-actin cloud, accounting for the DNA damage response signature at the gene level.

To test this idea, we used light sheet microscopy to measure the extent of nuclear compression in WT compared to *Dock8* KO T cells migrating under agarose, hypothesizing that the central F-actin in WT T cells plays a role in protecting the nucleus from force-mediated deformation. Indeed, we found the nuclei of *Dock8* KO T cells were significantly more compressed, and thus had a decreased height under agar than did the nuclei in WT T cells (**Fig. 5g,h**). To connect this observation with the gene expression results, we then asked whether the greater nucleus compression led to DNA damage by allowing WT and *Dock8* KO T cells to migrate in collagen for 12 hours and performing an alkaline comet assay, a method to detect double-stranded DNA breaks. Even with the exclusion of likely apoptotic cells (score 4, comparison with etoposide control which is a DNA damaging agent), we found a substantial increase in the frequency of cells with DNA breaks (scores 1-3) in migrating *Dock8* KO T cells (∼38%) compared to WT T cells (∼21%) (**Fig. 5i**). Taken together, our data suggest that Dock8 has a nucleo- protective role in migrating T cells, maintaining nuclear shape integrity and thus reducing mechanical force-induced genomic stress.

### Mst1 is required for the nucleo-protective central actin pool during migration

Interestingly, individuals with mutations in the serine-threonine protein kinase 4 (*Stk4*) gene, encoding for the mammalian sterile 20-like (Mst1) protein, present with an immunodeficiency that, while not entirely overlapping in clinical presentation, is reminiscent of Dock8-deficiency^50^. Like for Dock8, loss of Mst1 results in T cell lymphopenia and recurrent cutaneous viral and bacterial infections in particular, as well as eczema and atopic dermatitis.^51–55^ Naïve *Stk4* KO T cells have a defect in egressing the thymus and in trafficking to secondary lymphoid organs, and display reduced motility in 2D migration assays as well as within the lymph node^56, 57^. A direct link between Mst1 and Dock8 was suggested by studies showing that Dock8 can bind Mst1 through its N-terminal region^58, 59^, and that Mst1 regulates Dock8 via phosphorylation of Mps one binder 1 (Mob1), a Hippo pathway scaffold protein that can then interact with and activate Dock8 to promote Rac1 activity^56^.

Thus, we next asked whether Mst1 was involved in the Dock8-dependent and confinement-induced redistribution of F-actin in activated T cells. We isolated T cells from *Stk4* KO mice, activated them *in vitro* as before and confined them under agarose. Strikingly, we observed that *Stk4* KO T cells had a similar complete abrogation of the central F-actin pool under confinement as *Dock8* KO T cells, while being indistinguishable from WT T cells when migrating in 2D without confinement (**Fig. 6a**). The loss of the mechanically regulated redistribution of F-actin in *Stk4* KO T cells also led to their entanglement and death in 3D collagen matrices (**Fig. 6b-d**). The phenotype of *Stk4* KO T cells that had lost cell shape cohesion was near-identical to that of *Dock8* KO T cells (**Fig. 6b**), and *Stk4* KO T cells also had increased lamin A/C protein expression, scaling with the density of collagen they were embedded in (**Fig. 6e**). This suggested that *Stk4* KO T cells were experiencing increased mechanical stress similar to *Dock8* KO T cells.

**Figure 6.**
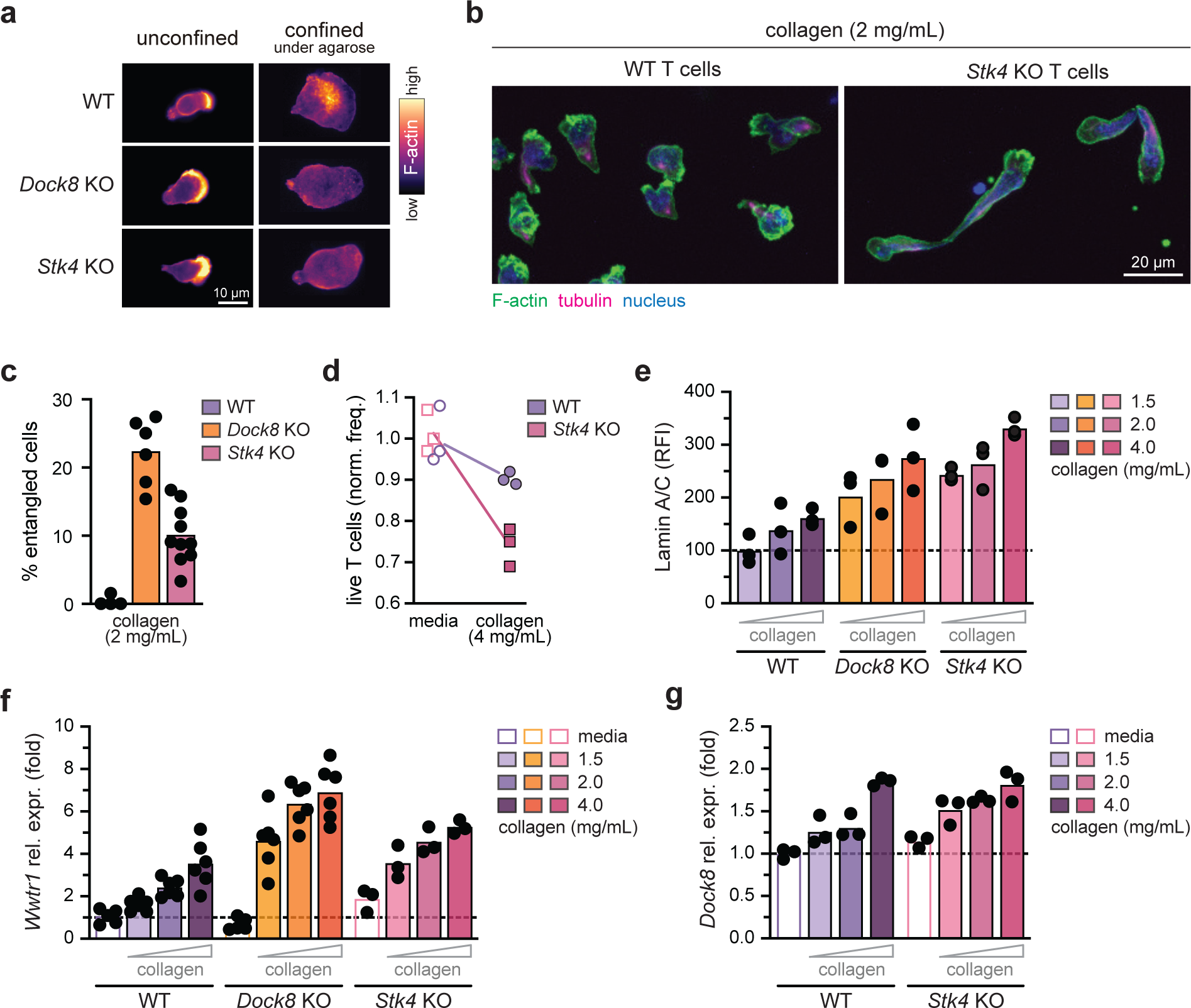
Mst1 is required for the nucleo-protective central actin pool during migration. (a) Fluorescence intensity of F-actin (phalloidin) in WT, *Dock8* KO, or *Stk4* KO T cells migrating on fibronectin-coated glass either in absence of (unconfined) or under 1.2% agarose (confined). Representative images (epifluorescence microscope) are shown. **(b)** Representative confocal microscopy images (max projection of z-stacks) of WT and *Stk4* KO T cells migrating through collagen (2 mg/ml). Fixed cells were stained for F-actin (phalloidin), tubulin (α-tubulin antibody), and nucleus (Hoechst). **(c)** Summary plot of percent WT, *Dock8* KO, and *Stk4* KO T cells entangled in collagen matrix (2 mg/ml) after 4 hours of migration. Data points, replicate T cell cultures from *n =* 4 mice per genotype; means are indicated. **(d)** Summary plot of the frequency of live WT or *Dock8* KO T cells either resting in media or after 24 hours of migration in 2mg/ml collagen matrices. Data summarized from *n =* 3 mice per genotype; frequency is normalized to the mean viability of WT T cells in media. (**e)** Lamin A/C expression quantified by flow cytometry in WT, *Dock8* KO, or Mst1 KO activated CD8^+^ T cells migrating in 1.5, 2, or 4 mg/ml collagen for 24 hours. Data summarized from *n =* 3 mice per genotype; lines, means; RFI is normalized to the mean of WT T cells in 1.5 mg/ml collagen. Dotted line at RFI = 100. **(f)** Gene expression of *Wwtr1* (Taz) assessed by quantitative RT-PCR in WT, *Dock*8 KO, or Mst1 KO activated CD8^+^ T cells cultured in media or migrating in 1.5, 2, or 4 mg/ml collagen for 24 hours. Expression was normalized to housekeeping gene Tbp and shown as fold change compared to WT T cells in media. Dotted line at fold = 1; data summarized from *n =* 3-6 mice per genotype from 1-2 independent experiment; lines, means. **(f)** Gene expression of *Dock8* assessed by quantitative RT-PCR in WT or Mst1 KO activated CD8^+^ T cells cultured in media or migrating in 1.5, 2, or 4 mg/ml collagen for 24 hours. Expression was normalized to a housekeeping gene and shown as fold change compared to the WT T cells in media. Dotted line at fold = 1; data summarized from *n =* 3 mice per genotype from 1 experiment; lines, means.

The Hippo pathway, of which Mst1/2 are core components in mammals, ultimately leads to the nuclear translocation of the transcriptional regulators Yap (yes-associated protein) and Taz (WW-domain-containing transcription regulator 1), encoded by *Yap* and *Wwtr1* respectively, which bind to DNA together with TEAD (transcriptional enhanced associated domain) transcription factors^60, 61^. Among other functions, Yap/Taz signaling has been shown to be critical in translating mechanical cues, including cell tension, ECM stiffness, cell density, and shear flow forces, into gene expression changes, including the upregulation of *Yap* and *Wwtr1* themselves^62^. To corroborate the hypothesis that the change in lamin A/C expression was a result of increased mechanosensing in *Dock8* KO and *Stk4* KO T cells, we measured *Wwtr1* expression by qPCR in cells incubated in media, or migrating in 1.5, 2, or 4mg/ml collagen for 24 hours. In WT T cells, *Wwtr1* expression was increasingly upregulated with greater collagen density compared to cells in media (**Fig. 6f**). Interestingly, the titrated response in *Wwtr1* was maintained in both *Dock8* KO and *Stk4* KO T cells, but the upregulation in *Wwtr1* was substantially greater even at low collagen densities compared to WT T cells (**Fig. 6f**). Given this result, we asked whether *Dock8* expression itself might also be mechanically regulated. Indeed, in both WT and *Stk4* KO T cells, *Dock8* transcripts increased with greater collagen density, also confirming that Dock8 expression was intact in Mst1-deficient cells (**Fig. 6g**). These data suggest that together, Mst1 and Dock8 are co-requisite for the mechanosensitive actin redistribution in T cells that is important for cell cohesion, survival, and nuclear protection during migration through confined spaces. Thus, *Dock8* and *Stk4*- deficiency results in cells experiencing greater mechanical forces, and responding with transcriptional changes accordingly, than WT T cells in the same environments.

## Discussion

In most physiological contexts, immune cell migration occurs in highly confined settings, constrained by tissue architecture, ECM structure, and cell density. Here, we report a mechanosensitive cytoskeletal response axis in migrating T cells and DCs. Our data show that Dock8, whose expression is restricted to hematopoietic cells, is required for the redistribution of F-actin under confinement, localizing actin polymerization to the cell center near the nucleus front. Moreover, we found that in addition to Dock8, F-actin rearrangement in confined migrating T cells is dependent on Mst1, a member of the Hippo signaling pathway known to transmit mechanical input into biochemical changes. Importantly, our data suggest that the redistribution of F-actin serves two important functions. One role of the F-actin relocalization is in the maintenance of cell shape integrity during the navigation of tight interstitial spaces. The second function of the central F-actin pool is in preventing DNA damage by shielding the nucleus from mechanical forces. Thus, we have identified a novel mechanism by which cells rapidly integrate mechanical cues as they move through environments that necessitate cell deformation, both in balancing speed with exploration, and in protecting the cell’s genomic material.

As leukocytes migrate in increasingly confined environments, the reduction of actin polymerization at the leading edge, with a concomitant increase in the cell center, appears to play a role in ensuring the maintenance of cell shape integrity while rapidly exploring interstitial space. The Arp2/3 and WAVE complex-dependent lamellipodial pool of F-actin is critical for cellular extrusion into the environment and hence promotes exploration^63, 64^, but this has to be balanced by actomyosin contractility in the cell rear^65, 66^. In *Dock8*-deficient T cells, increased actin polymerization at the lamellipodia results in the leading edge moving faster than the cell rear. This explains both why *Dock8* KO cells become elongated in collagen matrices, and why *Dock8* KO cells attain higher speeds in simple channels and pillar forests. When *Dock8* KO cells are navigating an obstacle while under confinement, they are unable to reconcile the opposing forces of two leading edges and the result is conflicting cell fronts that can ultimately rip cells apart. The role of Dock8 in reining in actin protrusions only becomes critical under confinement, because in obstacle-free settings the cell rear is able to ‘keep up’ with the cell front. This is in line with prior studies which establish that rear contraction is dispensable in less constrained environments^10^. Intriguingly, we found that unlike T cells and DCs, neutrophils did not exhibit the same F-actin rearrangement in response to mechanical pressure, and *Dock8* KO neutrophils did not lose cell cohesion while migrating in confined environments. One major difference between neutrophils and other leukocytes is their unique multi-lobulated nucleus, which allows for greater deformability^67^. Perhaps neutrophils, with a more malleable nucleus, and a reduced need to protect genomic material due to their terminal differentiation and shorter life span^68^, do not have the same requirement for a perinuclear F-actin structure when confined. Different immune cell types may thus have a differential dependence on Dock8 expression, and other mechanosensitive mechanisms by which neutrophils maintain cell cohesion during 3D migration independent of Dock8 will have to be further investigated.

The precise molecular machinery by which Dock8 relocalizes F-actin remains to be elucidated. Dock8 contains a DHR-1 domain, which binds PI(3,4,5)P3 (phosphatidylinositol 3,4,5- trisphosphate) in the membrane^69^. One hypothesis is that Dock8 actin regulation is mediated by PIP(3,4,5)P3, a lipid involved in promoting actin polymerization at specific regions of the cellular cortex^70^. Alternatively, the interaction of Dock8 with Cdc42 via its DHR-2 domain, which has specific catalytic activity as a GEF^25^, could play a role. In T cells, Dock8 exists in complex with WIP and WASp to promote Cdc42 activity^71^. In migrating DCs in 2D, loss of Cdc42 leads to the presence of multiple leading edges, and in 3D their motility is almost completely abrogated^72^. Interestingly, in T cells loss of Cdc42 or PAK1/2 led to a similar cellular elongation and loss of shape integrity as observed in Dock8-deficiency, but neither inhibition of Rac1 or WASp impacted cell shape during migration^13^. Dock8 interaction with WASp is therefore unlikely the mechanism of cytoskeletal coordination observed under confinement, but the localization or activation of Cdc42 within the cell may contribute. It was recently shown that WASp regulates the formation of small mechanosensitive actin foci which push outwards against the environment orthogonally to cellular movement^13^, and using TIRF microscopy we observed a reduction, albeit not a complete abrogation, of such foci in Dock8-deficient T cells. Thus, to what extent the central F-actin pool we describe, and the WASp-dependent actin foci that are also a response to compressive forces, are mediated by the same cytoskeletal circuitry remains to be defined.

Concurrent with the loss of cell cohesion in *Dock8* KO cells was the loss of nuclear shape integrity. Given our findings that *Dock8*-deficient T cells passed through narrow constrictions at both higher rates and faster passage times than WT T cells, it is possible that the central F-actin pool limits entry into very small constrictions. This is in contrast with an Arp2/3-dependent perinuclear actin pool that was found to transiently appear during DC passage through constrictions hypothesized to facilitate passage of the rigid nucleus^19^. A key difference could be the relatively softer nucleus of T cells, at least in part due to much lower lamin A/C levels which are an important determinant of nucleus rigidity^47, 73^. Importantly, whereas nuclear envelope rupture has been documented in DCs and cancer cells^41, 42^, it was unknown whether nuclear envelope ruptures occur in migrating T cells. Here we showed that such ruptures were rare in both WT and *Dock8* KO T cells, and that even in severely entangled *Dock8* KO cells the nuclear envelope remained intact. Indeed, mechanical strain has been shown to induce DNA damage also in absence of nuclear envelope rupture^74^, in line with the greater number of DNA breaks we observed in migrating *Dock8* KO T cells using a comet assay. That the *Dock8* KO T cells experience greater mechanical forces was also supported by the much more flattened nucleus under confinement compared to WT T cells, as well as the changes in gene expression observed only in migrating *Dock8* KO T cells, at least some of which are likely responses to greater DNA damage on the one hand, or increased mechanosensing on the other. Indeed, one read-out of the increased mechanosensitivity of *Dock8* KO cells was lamin A/C expression itself, which was higher in the KO cells both at the mRNA and the protein level compared to WT T cells. Lamin A/C expression has been shown to be upregulated in response to mechanical force, and this upregulation plays a protective role in mesenchymal cells, macrophages, and others^48, 49, 75^. In *Dock8* KO T cells, upregulation of lamin A/C was insufficient to prevent force-mediated DNA damage, but the phosphorylation state of lamins and other nuclear envelope proteins including emerin^76^, which we found was downregulated at the transcript level in KO T cells, may also play a role. Finally, consistent with an increase in mechanical force experienced by *Dock8* or *Stk4* KO T cells, we also detected an upregulation of *Wwtr1* expression with increased collagen density that was greater in both *Dock8* and *Stk4* KO T cells.

It is worth noting that while *Dock8* or *Stk4* KO T cells had a complete loss of the central actin structure under confinement in all cells examined, only a fraction of KO T cells became entangled in collagen at any given time. We do not address here whether there is heterogeneity between cells that contributes to differences in survival, but given that cell death increases as a function of collagen density, it is likely that chance encounters with specific environmental constraints account for at least some of the variability in loss of cell cohesion. To this point, we observed that in under agarose assays, cell confinement was not sufficient to elicit entanglement, but when cells encountered obstacles such as other cells or debris, this led to a stretched cell phenotype. This aspect of Dock8-dependent F-actin regulation may relate to the clinical phenotypes observed, as cellular survival defects in *Dock8*-deficiency have been described in T cells^24, 77, 78^, B cells^79^, NKT cells^80^, innate lymphoid cells^81, 82^, and DCs^23^. It is possible that in addition to migration-induced shattering of cell, sustained DNA damage resulting from nuclear stress during migration may also play a role in reducing cell viability over time. While in Mst1 immunodeficiency, contributors to the lymphopenia observed have been described to be severely reduced thymic egress and increased FAS-mediated apoptosis^51^, we describe for the first time that Mst1 is also required for cell cohesion during 3D migration via a mechanosensitive regulatory network with Dock8 and actin, which may explain some of the overlapping features of Mst1 and Dock8 immunodeficiencies.

The mechanobiology of immune cells is a rapidly developing field, and previous to this work, most of our understanding of T cell mechanosensing was with respect to synapse formation during priming or target cell killing. Much less is known about whether and how T cells, or immune cells more generally, adapt to confined 3D interstitial migration and to what extent mechanosensing in 2D compared to 3D differ^16, 83^. It was recently shown that mechanical input leads to transcriptional changes in T cells^84–86^. Moreover, stiffness sensing by effector T cells is a critical driver of the differentiation of tissue resident memory T cells, and confinement-induced motility plays a role in antigen surveillance by T cells in the salivary gland^44, 85^. Overall, defining the signaling axes which impact the ability of T cells to migrate through and establish residence in confined tissue environments may provide a therapeutic opportunity to modulate the migration to, and survival within, tissues where T cell surveillance is undesirable, for instance in autoimmunity. Alternatively, this work may ultimately enable the design of antigen-specific T cells for therapeutic settings where withstanding large mechanical forces is key to their effector function, as might be the case in anti-tumor responses. Overall, this work highlights how studying instances where cell migration goes wrong can shed light on the molecular mechanisms at play in effective migration through tissues.

## Materials and Methods

### Human samples

Participants were enrolled and informed consent obtained at the Center for Innovative Medicine (CIM), Research Institute of the McGill University Health Center (RI-MUHC), as approved by the McGill University Health Center Research Ethics Board (human ethics protocol #2023-8829). Volunteers included both males and females, in the age range of 21 - 45 years, and were sampled by venipuncture.

### Mice

C57BL/6J, B6 CD45.1 (jax #002014), and NLS-nTnG (jax #023537)^87^ mice were obtained from The Jackson Laboratory. LifeAct-GFP^88^ mice were kindly shared by Janis Burkhardt (University of Pennsylvania). *Dock8* KO mice were generated by genOway and shared by Helen Su (NIH)^23^. *Stk4* KO mice^89^ were shared by Dae-Sik Lim (Korea Advanced Institute of Science and Technology). *Tln1*^flox/flox^ mice^90^ were shared by Irah King (McGill University). All mouse strains were bred in-house and studies were performed in accordance with the Guide for the Care and Use of Laboratory Animals, with approval by the McGill University Facility Animal Care Committee. Mice were housed in the CMARC animal facility at McGill University at a temperature of 18- 24°C, 30-70% humidity, and 12h/12h light-dark cycles. Mice were used for experiments at 6-12 weeks of age. Both male and female mice were used and sex-matched where possible.

### Cell isolation and culture

<u>Murine T cells:</u> Inguinal, axillary, brachial, and cervical lymph nodes were harvested and crushed through a 70 µm filter. T cells were isolated with the EasySep Mouse Total T Cell Isolation Kit (StemCell) following the manufacturer instructions. Isolated T cells were resuspended in RPMI 1640 supplemented with 10% heat-inactivated FBS, 2 mM L-glutamine, 100 U/mL penicillin, 100 ng/mL streptomycin, 1 mM sodium pyruvate, 10 mM HEPES, 1% non-essential amino acids, and 5 µM 2-mercaptoethanol (complete RPMI) at 2.5×10^6^ cells/mL. For activation, T cells were supplemented with 2 µg/mL anti-CD28 (37.51; Biolegend), seeded in a F-bottom 96-well plate precoated with 3 µg/mL anti-CD3 (145-2C11; Biolegend) using 5×10^5^ cells (200 µl) per well, and incubated at 37°C, 5% CO2. Two days after seeding, cells were centrifuged at 300 *g* for 5 min at 4°C and cultured in flasks at 10^6^ cells/mL in complete RMPI with 20 ng/mL recombinant mouse IL-2 (rmIL-2; Biolegend). To generate talin-deficient T cells, T cells were incubated with 2 µM TAT-CRE Recombinase (Sigma) for 1 h, to excise the transgene flanked by *loxP* sites, before stimulation with rmIL-2. Activated T cells were used for experiments on day 4.

<u>Human T cells</u>: Lymphocytes were isolated from whole blood collected in sodium heparin tubes using Ficoll-Paque PLUS Density Gradient Media (Cytiva) following manufacturer instructions. CD8^+^ T cells were isolated with the EasySep Human CD8^+^ T Cell Isolation Kit (StemCell) using the manufacturer protocol. Isolated CD8^+^ T cells were resuspended at 2×10^6^ cells/mL in RPMI 1640 supplemented with 10% heat-inactivated FBS, 2 mM L-glutamine, 100 U/mL penicillin, 100 ng/mL streptomycin, 1 mM sodium pyruvate, 10 mM HEPES, 1% non-essential amino acids, and 5 µM 2-mercaptoethanol (complete RPMI). For activation, cells were supplemented with 25 µl/mL ImmunoCult Human CD3/CD28 T Cell Activator (StemCell) and 20 ng/mL Recombinant Human IL-2 (rhIL-2; Biolegend), seeded in a F-bottom 96-well plate using 4×10^5^ cells (200 µl) per well, and incubated at 37°C, 5% CO2. Complete RPMI supplemented rhIL-2 was added to the cells to reach a concentration of 2×10^6^ cells/mL on day 3 and 6 post-activation. Activated T cells were used for experiments on day 8.

<u>Murine BMDC:</u> Bone marrow from femurs and tibias was harvested by flushing bones with RPMI and passed through a 70 µm filter. Isolated bone marrow cells were resuspended in complete RPMI at 2.66×10^5^ cells/mL. For differentiation, cells were supplemented with 20 ng/ml GM-CSF (Biolegend), seeded into 6-well plates using 8×10^5^ cells (3 mL) per well, and incubated at 37°C, 5% CO2. 3 days later, 3 mL of complete RPMI supplemented with 20 ng/ml GM-CSF was gently added to each well. 3 days later, 3 mL of media was gently replaced from each well with fresh complete RPMI supplemented with 20 ng/ml GM-CSF. 2 days later, on day 8, the supernatant containing the BMDCs was harvested, centrifuged at 300 *g* for 5 min at 4°C, and the cells were resuspended in complete RPMI supplemented with 10 ng/ml GM-CSF at 10^7^ cells/mL. BMDCs were pulsed with 1 µg/ml LPS (InvivoGen) for 30 minutes, then washed with media prior to use in migration assays.

<u>Murine neutrophils:</u> Bone marrow from femurs and tibias was harvested by flushing bones with RPMI and passed through a 70 µm filter. Neutrophils were isolated from bone marrow cells using the EasySep Mouse Neutrophil Enrichment Kit (StemCell) following manufacturer instructions. Neutrophils were resuspended in complete RPMI supplemented with 20 ng/ml GM-CSF at 1×10^7^ cells/mL. Neutrophils were pulsed with 1 µg/ml LPS (InvivoGen) for 30 minutes, then washed with media prior to use in migration assays.

### Flow cytometry

Cells were distributed in a U-bottom 96-well plate prior to centrifugation at 300 *g* for 5 min at 4°C. Cells were stained in FACS buffer (PBS supplemented with 2% FBS and 5 mM EDTA) for 30 min at 4°C using the following antibodies: eFluor450 anti-mouse CD4 (GK1.5; Invitrogen), BV605 anti-mouse CD8 (53-6.7; Biolegend), anti-mouse TCR-β BV711 (H57-597; Biolegend), PE anti-mouse CD44 (IM7; Biolegend), and the Fixable Viability Dye eFluor 780 (eBioscience) for the exclusion of dead cells. After extracellular staining, the cells were fixed in 4% PFA in PBS at 4°C for 15 minutes. For the intracellular staining, cells were stained and permeabilized in FACS buffer supplemented with 0.3% Triton X-100 at 4°C for 1 hour using the following antibodies/probes: FITC or PE anti-mouse/human Lamin A/C (4C11; Cell Signaling Technology), AlexaFluor647 phalloidin (Invitrogen). Flow cytometry was performed on an LSR Fortessa (BD Biosciences). Files were analyzed using FlowJo (BD Biosciences).

### RNA extraction for sequencing and RT-qPCR

<u>RNA extraction</u>: RNA extraction was performed using the PureLink RNA Mini Kit (Invitrogen). 2 to 4×10^6^ CD8^+^ T cells were lysed in the lysis buffer provided and passed through a homogenizer (Invitrogen) before RNA purification following manufacturer instructions. RNA concentration was quantified using a NanoDrop (Thermo Scientific). Isolated RNA was stored at -80°C (for bulk RNA sequencing) or immediately converted to cDNA using the High-Capacity cDNA Reverse Transcription Kit (Applied Biosystems) following manufacturer instructions, using 1000 ng RNA per 20 µL mix and stored at -20°C until used.

Two-step RT-qPCR: qPCR was performed on a StepOnePlus Real-Time PCR System (Applied Biosystems) using 2 µL of cDNA per well combined with TaqMan Fast Advanced Master Mix, the TaqMan probe Mm01277042_m1 (*Tbp*) in VIC (endogenous control), and one of the following TaqMan probes, in FAM: Mm00613802_m1 (*Dock8*), Mm01289583_m1 (*Wwtr1*) (all reagents from Applied Biosystems).

### RNA sequencing

Purified RNA was provided to the Institut de recherches cliniques de Montréal sequencing core facility (QC, Canada). RNA quality control was performed by Pico assay with a bioanalyzer (Agilent). RNA-seq libraries were prepared from 150 ng RNA per condition using the following kits: RiboCop rRNA Depletion Kit HMR (Lexogen), KAPA RNA HyperPrep Kit (Roche), and TruSeq DNA UDI 96 Indexes (Illumina). Libraries were sequenced on a NovaSeq 6000 System (Illumina) using a Flow Cell S2 (Illumina) and a paired end run. More than 60 million paired-end reads were generated per sample. Reads were provided by the sequencing core facility as FASTQ files, quality checked using FastQC and MultiQC, and mapped to GRCm38/mm10 *Mus musculus* genome using HISAT2. Alignment quality checks were performed using Samtools Flagstat, FastQC, Picard Tools, and MultiQC. Sequencing reads overlapping exons were counted using featureCounts. Differential gene expression analysis was performed using EdgeR in R project^91^, filtering out low expression genes using the filterByExpr function, and normalizing the read counts (calcNormFactors function). The p values were corrected using the Benjamini-Hochberg method and only genes with an FDR < 0.01 were considered statistically significant. No fold change threshold was set unless stated otherwise. Volcano plots were generated using the ggplot2 package in R project. Heatmaps were built from CPM values using the R pheatmap function. CPM values were centered and scaled in the row direction. Gene ontology analysis was performed using Metascape^92^. Only genes that were significantly upregulated (FDR < 0.01 and fold change > 2) were included. Enriched clusters identified with less than 10 genes were filtered out from the analysis.

### Comet assay

An alkaline comet assay (capturing both single and double stranded DNA breaks) was performed using the Comet Assay Kit (Abcam) following the manufacturers protocol. Prior to the assay, a sample of activated T cells were treated with 10 µM etoposide for 1 hour at 37°C to generate a positive control. The comet tails were captured by epifluorescence and resulting images were manually scored as indicated in the figure.

### Collagen gel migration assay

<u>Gel preparation</u>: Collagen gels were prepared using type 1 bovine atelocollagen (Nutragen; Advanced Biomatrix) at concentrations of 1.5, 2, and 4 mg/mL as indicated in the figure legends. Collagen gels were prepared and mixed on ice in 12-well plates by sequentially mixing water, neutralization buffer, collagen, concentrated RPMI and cells (in FBS) to reach a final concentration 1X RPMI, 2 g/L NaHCO3, 25 mM HEPES, 2 mM L-glutamine, 20 ng/mL rmIL-2, and 12.5% FBS, pH = 7.2. Neutralization buffer consisted in 0.13N NaOH and was added at a ratio of 1 µl per 10 µl Nutragen. Concentrated RPMI consisted of 10X RPMI supplemented with 14 g/mL NaHCO3, 175 mM HEPES, 14 mM L-glutamine, 140 ng/mL rmIL-2, and NaOH to reach pH = 7.2. Cells were seeded at a final concentration of 2×10^6^ cells/mL. Gels were allowed to polymerize at 37°C, 5% CO2 for at least 2 h before individual assays. Gels were either fixed for staining or digested to extract the cells for downstream applications such as flow cytometry or RNA extraction.

Fluorescently labeled collagen: Fluorescent type I bovine atelocollagen was prepared by labelling Nutragen (Advanced Biomatrix) with Alexa Fluor 647 following a previously published protocol for labelling rat tail type I collagen^93^, with the following modification: the collagen gel was prepared using the recipe above without L-glutamine, rmIL-2 and FBS instead of the protocol recipe using DMEM and RB.

<u>Gel digestion</u>: Collagen gels were dissociated using pipette tips and digested at 37°C for 45 minutes using 2 mg/mL Collagenase D (Sigma) in combination with gentle agitation. Cells were collected in microtubes and centrifuged at 300 *g* for 5 min.

<u>Gel fixation and staining</u>: Collagen gels were submerged in ice-cold 4% PFA in PBS for 30 minutes at room temperature and then cells permeabilized using 0.5% Triton-X 100 in PBS for 30 minutes at room temperature. Gels were stained overnight in PBS supplemented with 2% BSA and 0.1% Triton-X 100 in PBS using the following antibodies/probes: Alexa Flour 647 phalloidin (Invitrogen), Hoechst 33342 or 34580 (Invitrogen), Alexa Fluor 488 anti-mouse/human ɑ-tubulin (B-5-1-2; Invitrogen). Collagen gels were rinsed with PBS prior to imaging by confocal microscopy.

### Under agarose migration assay

<u>Agarose preparation</u>: The under agarose migration assay was adapted from^43^. In brief, 35 mm glass-bottom dishes (Ibidi) were functionalized using a plasma cleaner (Harrick) and coated with 10 µg/ml fibronectin (Sigma) for 30 minutes at 37°C. Agarose gels were prepared by combining a solution made of 9 mL RPMI without phenol red, 10 µl 7.5% NaHCO3, 1 mL 10X HBSS, 1 mL FBS, and 20 µl 50 mM ascorbic acid, maintained at 56°C, to a solution of agarose made of 0.1 to 0.4 g of UltraPure Agarose (Invitrogen) melted in 9 mL of sterile water. A concentration of 1.2% agarose was used unless otherwise indicated. Fibronectin-coated dishes were rinsed with PBS, and 3 mL of agarose gel were poured per dish. Once solidified, agarose gel loading ports were made using a 2 mm UniCore punch spaced 2 mm apart. Cells were seeded in the first access port, at a concentration of 10^6^ to 10^7^ cells/mL in complete RPMI without phenol red. CCL19 was added to the second access port, at a concentration of 2 µg/ml. Cells were incubated at least 2 hours at 37°C and 5% CO2 prior to performing live imaging by epifluorescence microscopy or fixing the cells for staining.

<u>Cell fixation and staining</u>: To fix the cells and retain their morphology, entry ports were flooded with 1 mL of ice-cold 4% PFA for exactly 5 min at room temperature before gently lifting off the agarose pad. Fixation was prolonged without the agarose pad for 10 min at room temperature. After fixation, samples were permeabilized with 0.2% Triton X-100 for 10 minutes at room temperature, then blocked with PBS supplemented with 1% BSA (blocking buffer). Cells were stained overnight at 4°C in blocking buffer using the following antibodies/probes: Alexa Fluor 488 or Alexa Fluor 647 phalloidin (Invitrogen), Hoechst 33342 or 34580 (Invitrogen), anti-ɑ-tubulin (B-5-1-2; Invitrogen), Alexa Fluor 488 anti-acetylated tubulin (6-11B-1; Sigma), Alexa Fluor 488 DNAse 1 (Invitrogen), Mitotracker Deep Red (Invitrogen), Wheat Germ Agglutinin Alexa Fluor 594 (Invitrogen), Alexa Flour 647 anti-vimentin (D21H3; Cell Signaling Technologies). Samples were rinsed with PBS and imaged by epifluorescence microscopy as described below.

### Microfluidic devices

Microchannels were prepared from PDMS as previously described^67, 94^ from custom-designed molds obtained from 4DCell. Prior to use for migration assays, microchannels were functionalized using a plasma cleaner (Harrick) and coated with 10 µg/ml fibronectin (Sigma) for 1 h at 37°C. Microchannels were rinsed multiple times with complete RPMI without phenol red prior to seeding the cells in the access ports at 10^8^ cells/mL. Cells were incubated for at least 2 h at 37°C and 5% CO2 before imaging their dynamic behavior by epifluorescence microscopy as described below.

### Microscopy

Epifluorescence microscopy was performed with a ZEISS Axio Observer Fully Automated Inverted Microscope equipped with a Plan-Apochromat 20×/0.8 NA objective for all assays, except for the comet assay where a Plan-Neofluar 10×/0.3 NA objective was used. Confocal microscopy was performed with a ZEISS LSM880 equipped with a Plan-Apochromat 20×/1.0 NA water immersion objective. Total internal reflection fluorescence (TIRF) microscopy was performed with a (TIRF)-Spinning Disk Spectral Diskovery System (Spectral Applied Research) coupled to a DMI6000B Leica microscope equipped with a Plan-Apochromat 63×/1.47 NA oil immersion DIC objective. TIRF imaging depth was set to 120 nm. Lattice light-sheet microscopy was performed using a Zeiss Lattice Lightsheet 7 equipped with a 13.3×/0.4 NA objective for illumination, a 44.83×/1.0 NA objective for detection, and using a Sinc3 100 × 1800 lattice light sheet for acquisition. All live imaging was performed at 37°C with 5% CO2 using a top-stage incubator (Live Cell Instrument).

### Image analyses

<u>Dynamic shape plots</u>: Dynamic shape plots were generated from maximum intensity z-projections of time-lapse microscopy image sequences converted to evenly spaced sub-stacks of 10 to 20 frames to visualize cell morphology changes on a 1-minute scale. Sub-stacks were binarized in Fiji using Triangle thresholding. Cell outlines were extracted from the binary image sequence using the polygon selection tool.

<u>Principle component analysis (PCA)</u>: T cells embedded in collagen gels were fixed and imaged in 3D by confocal microscopy. Cells were segmented using the Surface tool in Imaris Image Visualization and Analysis software v. 10.1.1 (Oxford Instruments). Surfaces corresponding to cell debris, diving cells, dead cells, and cells partially contained in the imaging area were excluded. Remaining surfaces were used to create a mask of viable single cells. Any (partially) overlapping cells were separated into different channels to ensure the most accurate shape analysis. 3D masked images were converted to 2D images by maximum intensity z-projection in Fiji. The 2D projections were binarized by *Triangle* thresholding. Cell shape parameters were extracted using the built-in Analyze Particles function and the MorphoLibJ plugin. A PCA was performed using the prcomp function from the R statistical computing package^91^. ggplot2 and factoextra packages were used for data visualization. PC1 > 5 was considered as a threshold above which cells were entangled/had lost cell cohesion in the collagen.

<u>Heat maps</u>: Actin distribution heat maps were generated from time-lapse widefield epifluorescence images, based on protocol outlined in^95^. Cells were binarized by Triangle thresholding in Fiji. 2D average intensity z-projections were generated for each cell to yield an average cell shape over time. The 2D projections of all cells for each condition were combined into a single image stack and normalized to ensure a 30%-pixel saturation in each image. To account for differences in size between cells, the combined image stack was scaled to the smallest cell in the sequence. 2D average intensity z-projections were generated from the combined image sequence for each condition.

<u>Centroid RFI</u>: Cell centroid measurements were extracted with Fiji from widefield images of phalloidin-stained T cells. Centroid points were manually defined as the intersection of one axis along the direction of migration, and its perpendicular axis. Non-polarized, clumped, or dead cells were excluded from the analysis.

<u>Intensity histograms</u>: Fluorescence intensity histograms were generated by manually defining a line along the axis of migration. Non-polarized, clumped, or dead cells were excluded from the analysis. Intensity values along the line were extracted using the line profile tool in Fiji. A Python script was used to normalize x-axis values and perform linear interpolations.

<u>Nuclear height measurement</u>: Nuclear heights were extracted from the NLS-nTnG signal of T cells migrating under agarose as acquired by lattice light sheet microscopy. Non-polarized, clumped, or dead cells were excluded from the analysis. The thickness of the center of the nucleus in the Z-direction was measured in Imaris Image Visualization and Analysis software v. 10.1.1 (Oxford Instruments).

### Statistics

For analyses, statistical differences between groups were evaluated by the two-sided tests reported in figure legends. Comparisons were considered significant when *P* ≤ 0.05. All statistical tests were performed using Prism 10 (GraphPad). *P*-values are indicated in figures as falling into one of 4 categories: ns, nonsignificant, \**P* < 0.05, \*\**P* < 0.01, or \*\*\**P* < 0.001, and statistical tests performed are indicated in legends.

### Data Availability

RNA sequencing dataset is available through GSExxx (currently under submission). All other data from this study are available from the corresponding author upon request.

## Acknowledgements

We thank all the volunteers who kindly donated blood samples for our experiments. We thank Karel Prud’homme, Genvieve Perreault, and CMARC for their excellent care of our mice. The work was made possible by the McGill University Flow Cytometry Core Facility (FCCF) and Advanced Bioimaging Facility (ABIF). We particularly thank Nelly Vuillemin (ABIF) for her lattice light sheet expertise, and Tanner Ducharme for custom Python scripts. We thank all past and present Mandl lab members, especially Maryl Harris, Dakota Rogers, Angela Mingarelli, and Sam Jamaleddine for their contributions, as well as Reza Sharif-Naeini (McGill University), Johannes Textor (Radboud University, The Netherlands), and Stephanie Eisenbarth (Northwestern University) for their feedback. We are especially grateful to Michael Sixt, Patricia Reis-Rodrigues, and Alba Juanes Garcia at IST Austria for all their input and collaborative spirit in discussions we have had over the years this work was done.

## Funding

Fonds de Recherche du Québec – Santé (FRQS) fellowship 2023-2024 - BF2 - 334876 (CSh) Canadian Institutes of Health Research (CIHR) fellowship 201911FBD-434608-297037 (CSh) Human Frontiers in Science Program (HFSP) Long-Term Fellowship LT000110/2019-L (JP) Human Frontiers in Science Program (HFSP) grant RGP0053/2020 (JNM) Canadian Institutes for Health Research project grants PJT-462910 (JNM) and PJT-178083 (JFC) Canada Research Chair (JNM, JFC) McGill University Mi4 seed grant #5 (AS and JNM)

## Author contributions

J.N.M. conceived and supervised the project. J.N.M. and C.Sh. designed the research. C.Sh. performed most of the experiments with critical help from J.P., M.M., D.P., A.B., V.L., A.A., and C.Sc. J.P. contributed the RNA sequencing experiment and analysis, and established the microfluidic assays. M.M. contributed the neutrophil migration data; A.C. performed the PCA analyses of cell shape parameters and generated the F-actin heatmap summaries. Data analysis was led by C.Sh. and J.P. with key contributions from A.C., M.M., D.P., and A.B. At the McGill University Health Centre, A.S. coordinated the human ethics protocol with help from D.P. and oversaw the collection of human blood samples. Critical reagents and intellectual input were provided by V.L., A.A., W-K.S, A.S., A.E., and J-F.C. Manuscript figures were put together by C.Sh., J.P., and J.N.M. The manuscript was written by J.N.M and C.Sh, with all authors contributing and providing feedback.

---

**Figure S1. *Dock8* KO T cells are chemotactic but lose cell cohesion in collagen matrices. (a,b,c)** Summary plots of mean track speed (a), track straightness (b), track displacement (c), and track displacement angle (c) for WT and *Dock8* KO T cell migrating through 2 mg/mL collagen matrix towards a chemotactic gradient of 2 µg/ml CCL19. Data is *n =* 278 (WT) and *n =* 242 (KO) cells. **(d)** Examples of WT and *Dock8* KO T cells migrating through decision points in 2 mg/mL collagen matrix. Fixed cells were stained for tubulin (α-tubulin antibody). **(e,f)** Collagen matrices at various densities (1.5, 2.0, and 4.0 mg/ml) were imaged by confocal microscopy (e) and elastic modulus was measured by nanoindentation (f). Data is *n* = 4 technical replicates; lines: means; error bars: SD. **(g)** Representative z-stack projections of WT and *Dock8* KO T cells migrating in varying collagen densities (1.5, 2, and 4 mg/ml). Fixed cells were stained for F-actin (phalloidin), tubulin (α-tubulin antibody), and nucleus (Hoechst). Statistical tests: Mann-Whitney tests (a,b); Two-way ANOVA with Tukey’s multiple comparison test (f). \*\*\**P <* 0.001; ns, non-significant.

**Figure S2. Perturbed nuclei of entangled *Dock8* KO T cells are stretched but do not rupture. (a)** Timelapse example of WT T cell passing through a 2.5 µm constriction with fluorescent nuclear reporter NLS-nTnG. Point of nuclear rupture is indicated by *. **(b)** Timelapse example of *Dock8* KO T cell expressing LifeAct-GFP and NLS-nTnG migrating in 2 mg/ml fluorescent collagen matrix acquired by lattice light sheet microscope and shown as a z-projection.

Figure S3. *Dock8* KO T cells display normal cell cytoskeleton, organelle organization, total F-actin, and cell division under confinement. (a-e) Examples of WT and *Dock8* KO T cells migrating under 1.2% agarose. Fixed cells were stained for nucleus (Hoechst) (a-e), acetyl- tubulin (ac-tubulin antibody) (a), tubulin (α-tubulin antibody) (a), F-actin (phalloidin) (b-d), G- actin (DNAseI) (b), mitochondria (Mitotracker) (c), lysosomes (wheat germ agglutinin) (d), and/or vimentin (vimentin antibody) with cell outline from brightfield (e). **(f,g)** Total cellular F-actin of activated T cells (TCR-β+ CD44+) embedded in 2mg/mL collagen for 2 hours prior to digestion and stained with phalloidin for flow cytometry. Representative histogram (f) and data summarized from *n =* 4 mice per genotype (g). **(h,i)** Examples of *Dock8* KO T cells under 1.2% agarose that are stretching (h) or dividing (i). Fixed cells were stained for acetyl-tubulin (ac-tubulin antibody), tubulin (α-tubulin antibody), and nucleus (Hoechst).

**Figure S4. Confinement-dependent central actin pool is present in dendritic cells, but not in entanglement-resistant neutrophils. (a-c)** Representative examples of WT and *Dock8* KO BMDCs (a), T cells (b), and neutrophils (c) migrating under 1.2% agarose with a chemokine gradient (CCL19 for BDMC and T cells, fMLF for neutrophils). Fixed cells were stained for F-actin (phalloidin) and the nucleus (Hoechst). **(d)** LifeAct-GFP and NLS-nTnG WT and *Dock8* KO neutrophils spontaneously migrating through a 2mg/ml bovine collagen matrix (no chemokine added). **(e)** Summary plot of neutrophil entanglement frequency during migration in collagen matrix (d) from *n =* 3 mice per genotype, 2 independent experiments.

## Supplemental tables

Table S1. List of summary cell shape parameters used in the PCA with their relative contribution to variance per column.

## Supplemental movies

**Video S1**. Examples of LifeAct-GFP and NLS-nTnG WT and *Dock8* KO T cells moving in collagen gels, corresponding to Figure 1a.

**Video S2**. Examples of LifeAct-GFP and NLS-nTnG *Dock8* KO T cells entangled in collagen matrices.

**Video S3**. Examples of LifeAct-GFP NLS-nTnG WT and *Dock8* KO T cells migrating in 6 µm straight microchannels, corresponding to Figure 2b. Scale bar is 10µm.

**Video S4**. Examples of LifeAct-GFP NLS-nTnG WT and *Dock8* KO T cells migrating in pillar forest microchannel, corresponding to Figure 2d. Scale bar is 20µm.

**Video S5**. Examples of passing and non-passing LifeAct-GFP NLS-nTnG WT T cells in constriction microchannels, corresponding to Figure 2f. Scale bar is 10µm.

**Video S6**. Example of nuclear rupture in LifeAct-GFP NLS-nTnG WT T cell during passage through constriction microchannel, corresponding to Figure S2a.

**Video S7**. Example of LifeAct-GFP NLS-nTnG *Dock8* KO T cell migrating through fluorescent collagen imaged by lattice light sheet, corresponding to Figure S2b.

**Video S8**. Examples of LifeAct-GFP WT and *Dock8* KO T cells migrating under agarose imaged by TIRF, corresponding to Figure 3c.

**Video S9**. Examples of LifeAct-GFP WT and *Dock8* KO T cells migrating under agarose imaged by widefield microscopy, corresponding to Figure 3d.

**Video S10**. Example of LifeAct-GFP NLS-nTnG WT T cell migrating under agarose imaged by widefield microscopy, corresponding to Figure 4d

## Notes

### Competing Interest Statement

The authors have declared no competing interest.

