## Supplementary figures and images for "DOCK8 regulates a mechanosensitive actin redistribution that maintains immune cell cohesion and protects the nucleus during migration"

### Supplemental Figures

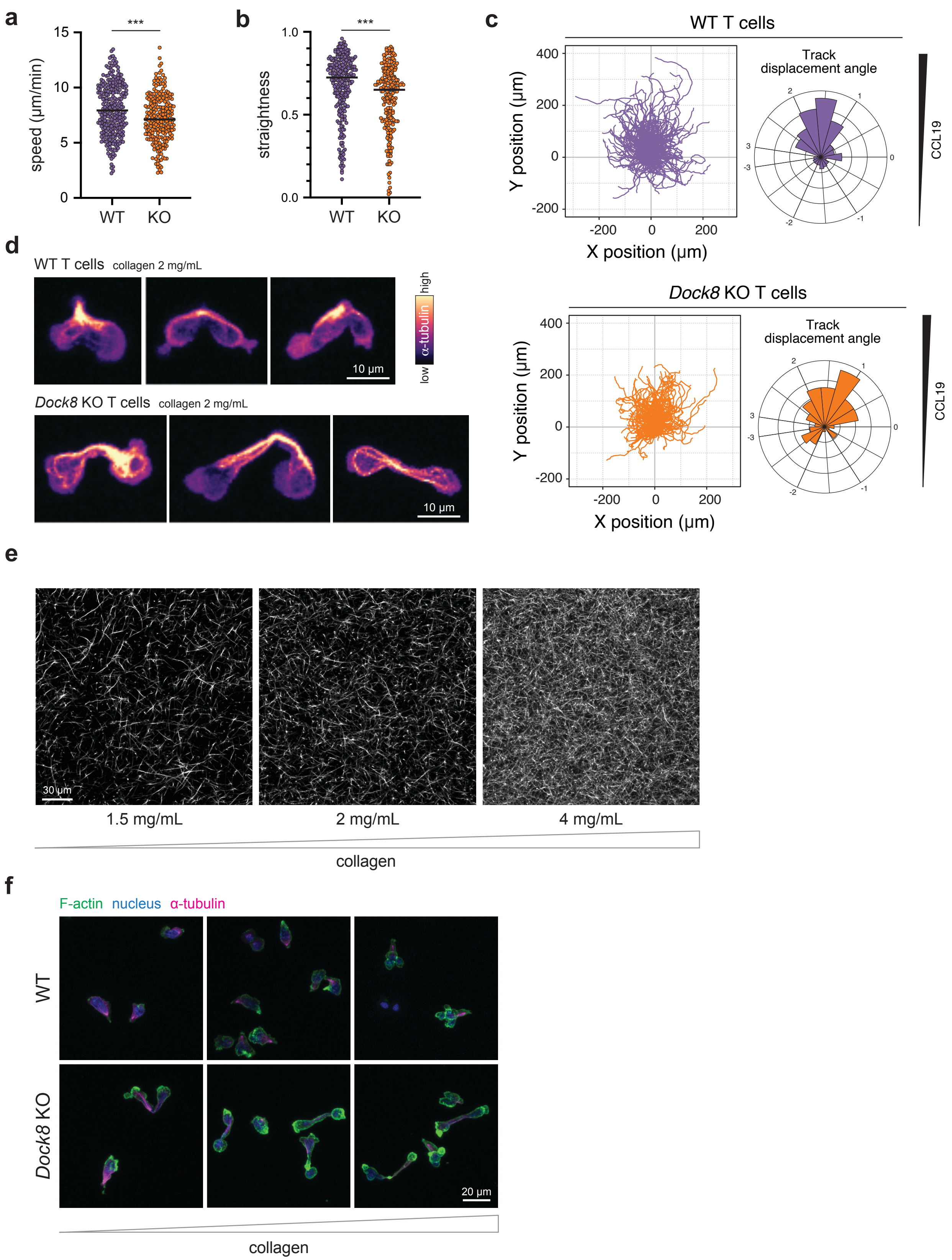

#### Shen *et al*_Figure S2

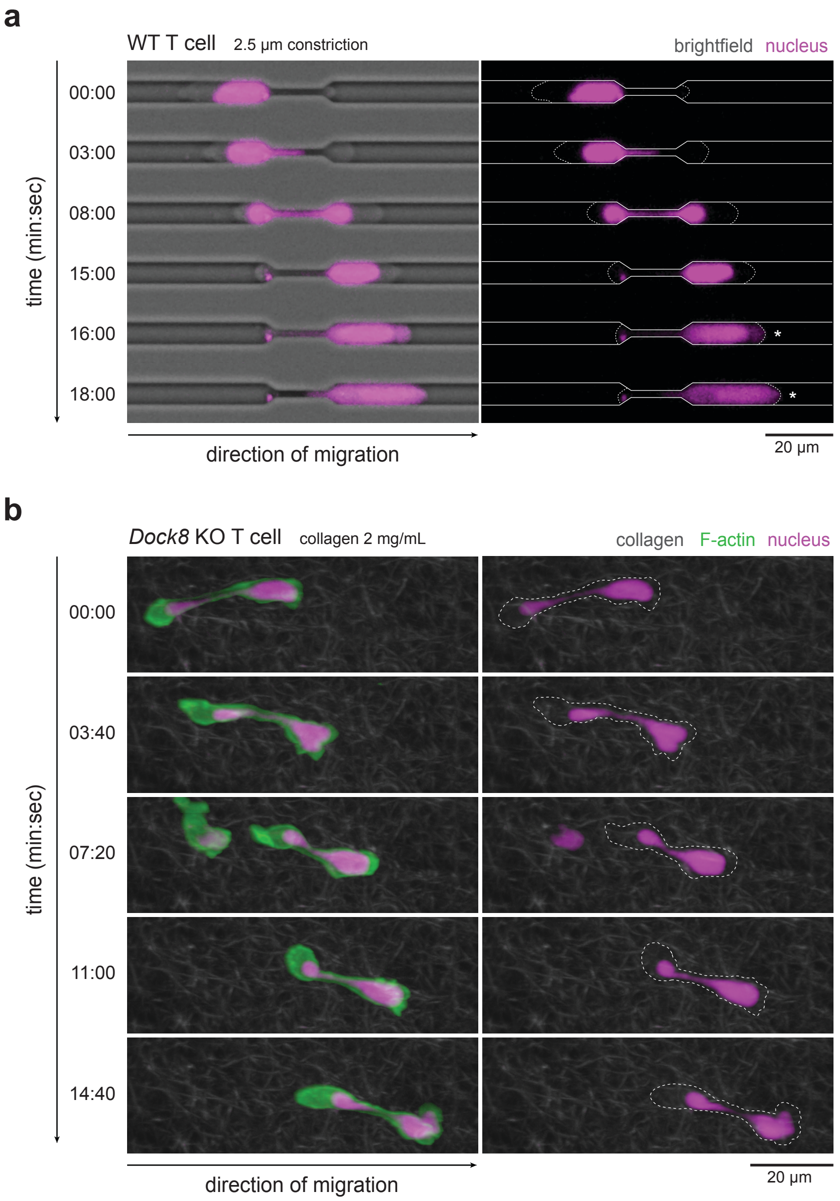

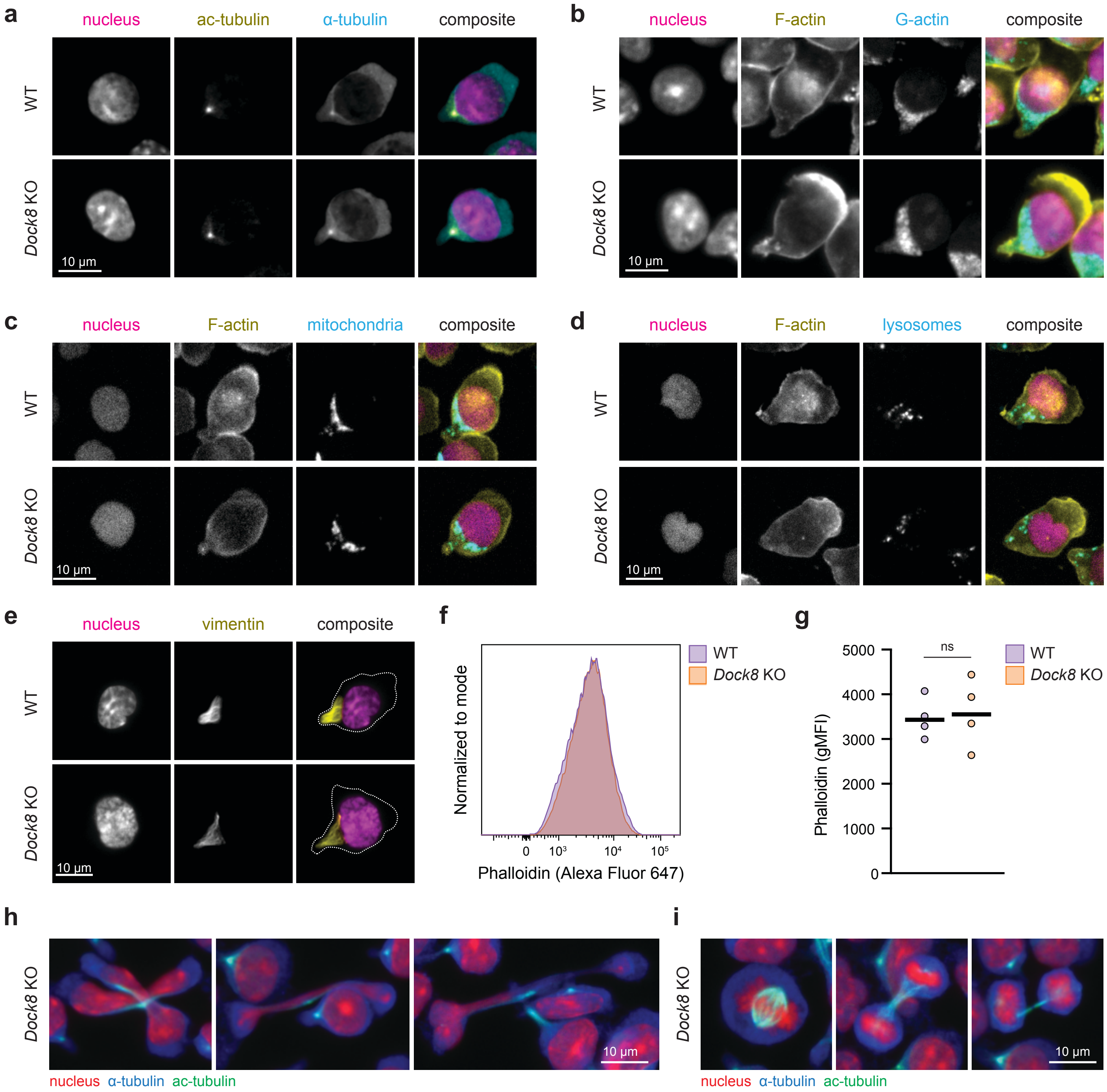

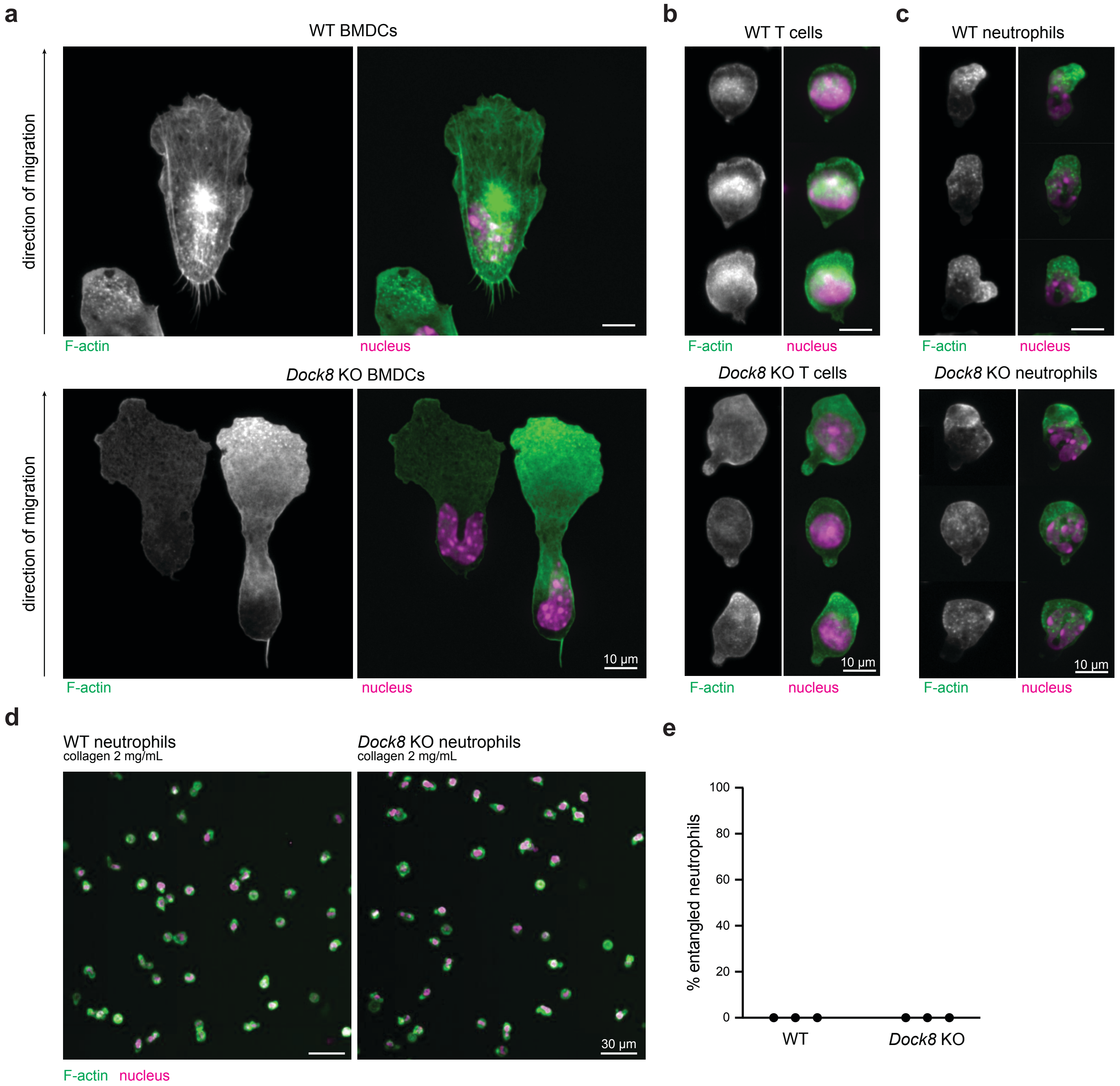
