## Supplemental Table 1 for "DOCK8 regulates a mechanosensitive actin redistribution that maintains immune cell cohesion and protects the nucleus during migration"

**Supplemental table 1. Summary cell shape parameters used in PCA**

| | PC1 | **PC2** |
|---|---|---|
| **Feret** | 7.72004300835294 | 0.00741839495603561 |
| **MinFeret** | 1.95767756546471 | 15.1825748079231 |
| **AR** | 4.84748793561407 | 5.59159166783884 |
| **Round** | 3.93509376733553 | 7.01458582234766 |
| **Solidity** | 7.0474781230942 | 0.175751614178413 |
| **Area** | 4.97538388527173 | 5.27952963677887 |
| **Perimeter** | 7.68927954352132 | 0.740475077218095 |
| **Circularity** | 6.19732071171204 | 1.32766424473589 |
| **Ellipse.Radius1** | 7.58797050592475 | 0.0015313455808298 |
| **Ellipse.Radius2** | 1.09913958610241 | 17.5945595074921 |
| **Ellipse.Elong** | 4.85411093567498 | 5.57733974887624 |
| **Convex.Area** | 6.5869972463315 | 2.72441495200489 |
| **Convexity** | 7.11539269384822 | 0.116553618892055 |
| **Obox.Length** | 7.68641791926519 | 8.37054047824699E-05 |
| **Obox.Width** | 1.95767756546471 | 15.1825748079231 |
| **GeodesicDiameter** | 7.77871172749818 | 0.074476434011451 |
| **Tortuosity** | 1.76939987453888 | 0.839894204958541 |
| **IncrDisc.Radius** | 0.238084266883591 | 13.5737718093963 |
| **AverageThickness** | 1.89532141744331 | 7.76646605954072 |
| **GeodesicElongation** | 7.06101172065773 | 1.2287425399421 |
